# Cloche/Npas4l is a pro-regenerative platelet factor during zebrafish heart regeneration

**DOI:** 10.1101/2025.02.19.639191

**Authors:** Junjie Hou, Yabing Song, Chenglu Xiao, Yuanyuan Sun, Jie Shen, Xiaokai Ma, Qinchao Zhou, Shih-Ching Chiu, Yang Xu, Yanyi Huang, Ye-Guang Chen, Xiaojun Zhu, Jianbin Wang, Jing-Wei Xiong

## Abstract

Zebrafish has full capacity of heart regeneration, but little is known about how blood cells, especially platelets, are involved in this regenerative process. Here we report that *cloche/npas4l* is a pro-regenerative platelet factor for heart regeneration. We found that haploinsufficiency of *npas4l* disrupted cardiomyocyte (CM) and endothelial cell (EC) proliferation and heart regeneration after injury. A single-cell transcriptomic atlas revealed that *npas4l* was dynamically expressed in platelets after heart injury and regulated robust interactions between platelets-CMs or -ECs. Decreasing platelets by either nitroreductase-mediated platelet depletion or platelet-deficient mutant *mpl* impaired CM and EC proliferation, and over-expression of *npas4l* in platelets sufficiently induced CM and EC proliferation in uninjured hearts, as well as rescued CM and EC proliferation defects in *cloche* mutants. Mechanistically, Npas4l positively regulated a panel of ligands expression including *bmp6* in platelets to fine-tune CM proliferation and heart regeneration. Therefore, this work unveils a novel platelet Npas4l signaling and presents mechanisms on how platelets modulate CM/EC proliferation *via* ligand-receptor network during zebrafish heart regeneration.

## Introduction

Organ regeneration remains a century-old question in spite of tremendous efforts to explore the replenishment of lost cells and tissues. In mammals, heart regeneration is restricted to before postnatal day 7, whereas some lower vertebrates like zebrafish keep this regenerative capacity over a lifetime (Tzahor and Poss, 2017; Zheng et al., 2021). Because of a lack of adult cardiac stem cells, it is generally believed that injury-induced de-differentiation and proliferation of CMs contributes to new myocyte regeneration (Jopling et al., 2010; Kikuchi et al., 2010; Ogawa et al., 2021; Xiao et al., 2016). In addition, the synchronized regeneration of coronary vessels, along with the precise modulation of cardiac fibrosis and inflammation, create a permissive environment with essential oxygen and nutrient supplies, thereby facilitating a regenerative niche for the proliferation of new myocytes. Over the past two decades, Neuregulin1 (Nrg1), yes-associated protein (Yap), miRNA-199a-3p, Krüppel-like factor 1 (Klf1), and the small-molecule cocktail 5SM are known to efficiently promote CM proliferation, likely *via* de-differentiation in zebrafish and mouse heart regeneration (Bersell et al., 2009; Du et al., 2022; Eulalio et al., 2012; Gemberling et al., 2015; Monroe et al., 2019; Ogawa et al., 2021). Other studies have shown that VEGF-Kdrl, Cxcl12b-Cxcr4a, and Vegfc-Vegfr3 are essential for cardiac angiogenesis and lympho-angiogenesis during zebrafish heart regeneration (Harrison et al., 2015; Harrison et al., 2019; Marin-Juez et al., 2016), and these signals are mainly secreted by CMs or ECs. However, injured hearts are composed of various cell types that coordinate to achieve this remarkable task in a spatiotemporal fashion. It remains incompletely understood how non-myocyte signals regulate CM and coronary vessel regeneration.

Derived from megakaryocytes, platelets are produced in the bone marrow with the characteristics of small size and no nuclei in mammals. Since platelets mediate thrombosis and hemostasis, they are widely involved in wound healing for bleeding arrest. However, due to their engagement in the thrombosis-inflammation response, platelets are recognized as crucial players in regulating the immune response by releasing cytokines and chemokines, thus creating an environment for tissue remodeling after injury (Mezger et al., 2019). Moreover, it has been shown that platelet activation modulates heart and brain repair by regulating tissue inflammation after ischemia-reperfusion injury (Rohlfing et al., 2022). Nevertheless, as an intricate secretory system, platelets release various growth factors, chemokines, and cytokines; however, their function in heart regeneration has never been addressed (Barrientos et al., 2008; Golebiewska and Poole, 2015; Grover et al., 2018; Versteeg et al., 2013).

The zebrafish *cloche* mutant was first discovered to affect both hematopoietic and EC lineages during zebrafish embryogenesis in 1995 (Stainier et al., 1995), the first genetic interval of *cloche* was defined at the telomeric region of chromosome 13 in 2008 (Xiong et al., 2008), and *npas4l*, a bHLH-PAS transcription factor, was cloned from *clo^s5^* in 2016 (Reischauer et al., 2016). As the earliest gene regulating the origin and formation of hemangioblasts, *npas4l* has mostly been investigated during zebrafish embryogenesis, but its function in organ regeneration especially heart regeneration has not been addressed. In this work, we aimed to search for critical molecular and cellular signaling on zebrafish heart regeneration by using single-cell RNA-sequencing (scRNA-seq) technology, *cloche* mutant, and heart injury models. We found, for the first time, that platelets and their Npas4l-Bmp6 signaling were essential for zebrafish heart regeneration, thus not only advancing our understanding of basic mechanisms on heart regeneration but also elucidating their potential applications for platelets and platelet-derived factors in cardiac repair and regeneration.

## Results

### Haploinsufficiency of *npas4l* disrupts zebrafish heart regeneration

In search of abnormal heart-regeneration phenotypes in heterozygous zebrafish mutants that are normally embryonic-lethal in homozygous mutants, we found that either *clo^m39/+^* or *clo^fv087b/+^ mutants* have abnormal heart regeneration (Reischauer et al., 2016; Xiong et al., 2008); *clo^m39^ is* a spontaneous deletion allele, while *clo^fv087b^* is an ethylnitrosourea-induced allele in which the mutation is unknown (Xiong et al., 2008). Based on the previous report that *cloche* encodes *npas4l* (Reischauer et al., 2016), we performed Sanger sequencing of the *npas4l* coding region of wild-type sibling and *clo^fv087b^* homozygous mutant embryos, and found that in the *clo^fv087b^* mutants, the transcription start site (ATG) of *npas4l* was mutated to AAG (Figure 1A), likely resulting in blockade of *npas4l* translation. Since it had been shown that *npas4l* expression was eliminated in *clo^m39^* mutants and either *clo^m39^ or clo^fv087b^* homozygous mutants are embryonic lethal (Reischauer et al., 2016; Stainier et al., 1995), we turned to assess *npas4l* mRNA expression in *clo^fv087b/+^* adult hearts by comparing with wild-type sibling hearts after ventricular resection, and found that *npas4l* mRNA in *clo^fv087b/+^* hearts decreased ∼50% (Figure 1B). By generating an anti-Npas4l antibody against a 253 carboxyl-terminal amino-acid peptide, we found that this antibody specifically recognized over-expressed Myc-tagged Npas4l proteins in 293T cells by western blot analysis (Figure S1C). Additionally, immunofluorescence staining revealed nuclear localization of Npas4l signals in conjunction with Myc-tag signals (Figure S1D). This antibody recognized endogenous Npas4l proteins in zebrafish hearts, revealing that the levels of Npas4l protein were lower in *clo^fv087b/+^*mutant hearts than wild-type sibling hearts at 1 day post-amputation (dpa) (Figures 1B and 1C). These data suggest that the *clo^fv087b/+^* mutant has *npas4l* haploinsufficiency with both reduced mRNA and protein expression. To elucidate the function of *npas4l* haploinsufficiency in heart regeneration, we assessed fibrosis resolution and cardiac cell proliferation in both *clo^fv087b/+^* and *clo^m39/+^* mutants after heart amputation (Figure 1D). By comparing wild-type sibling and *clo^fv087b/+^* mutant hearts, we categorized the hearts into three levels of regeneration, not regenerated (severe), partially regenerated (partial), and fully regenerated (regenerated), based on the serial-section acid-fuchsin orange-G (AFOG) staining (Figure S1A). We found the *clo^fv087b/+^*mutant exhibited more defective cardiac regeneration indicated by more fibrotic scars with AFOG staining and decreased myocardial regeneration labeled with myosin heavy chain (MHC) immunofluorescence (Figures 1E and S1A). Consistent with this, we found that *clo^m39/+^*hearts also had more fibrosis and collagen deposition and less regenerated myocardium than wild-type hearts at 30 dpa (Figure 1F). Thus, *npas4l* haploinsufficiency impairs zebrafish heart regeneration.

**Figure 1.**
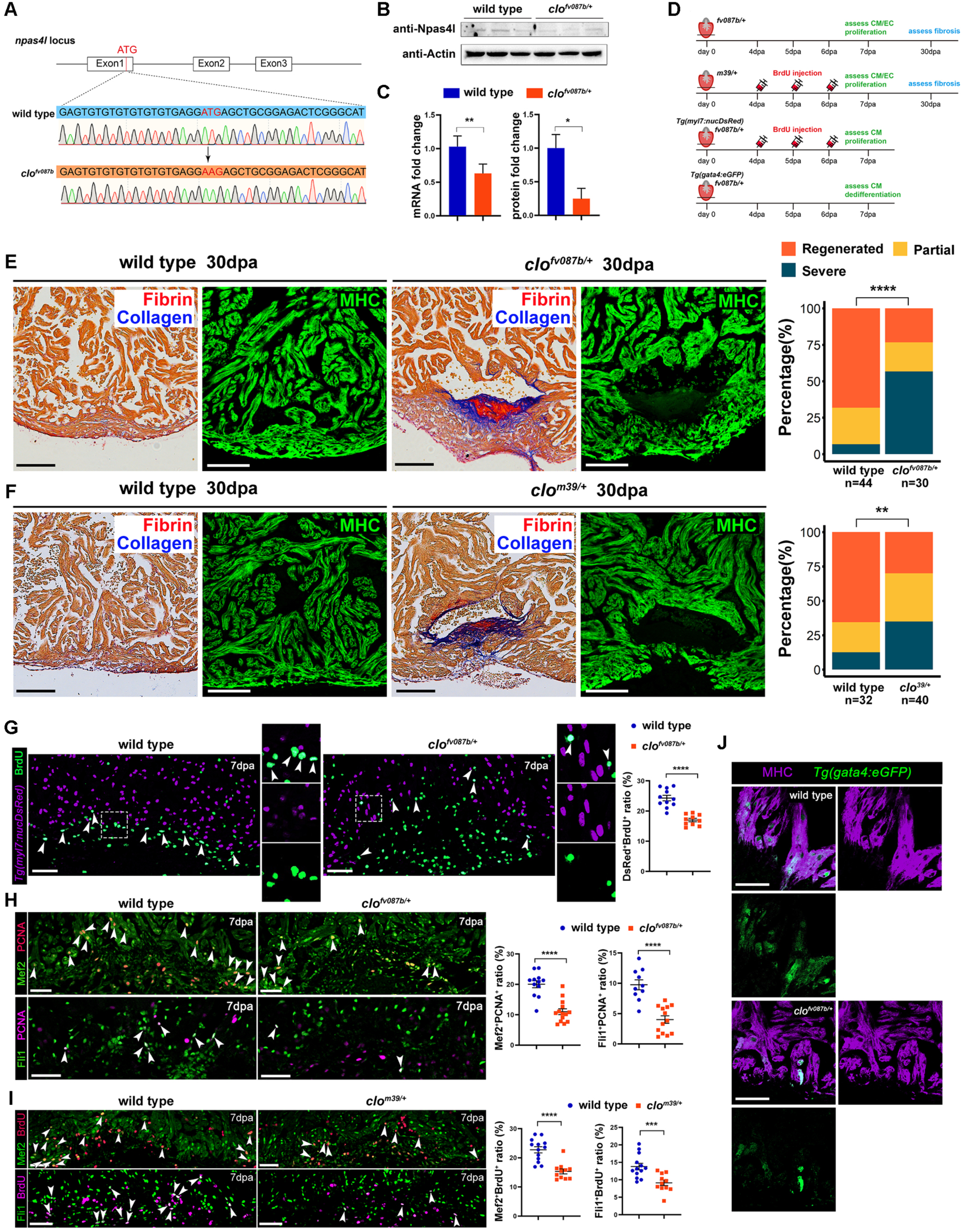
Haploinsufficiency of *npas4l* disrupts zebrafish heart regeneration. **A** Sanger sequencing of the *npas4l* locus of genomic DNA reveals that ATG in wild-type embryos (n=5) is mutated to AAG in *clo^fv087b^*mutant embryos (n=6) at 4 days post fertilization. **B, C** Quantitative RT-PCR and protein quantification (**C**) and western blot (**B**) of *npas4l* expression in adult zebrafish hearts at 1 dpa. mRNA loading is normalized by *gapdh* and protein loading by β-actin. n=3 replicates per group, and 10 hearts pooled together per replicate. Data are the mean ± SEM.; *p <0.05, **p <0.01; unpaired, two-tailed Student’s *t* test. **D** Schematic diagram of experimental design to explore *npas4l* function. **E** Representative images and quantification of AFOG and MHC immunofluorescence staining in wild-type sibling (n=44) and *clo^fv087b/+^* (n=30) hearts at 30 dpa. Data represents the percentage of each group; ****p <0.001; Pearson chi-square test. Scale bars, 100 μm. **F** Representative images and quantification of AFOG and MHC immunofluorescence staining in wild-type sibling (n=32) and *clo^m39/+^* (n=40) hearts at 30 dpa. Data represents the percentage of each group; **p <0.01; Pearson chi-square test. Scale bars, 100 μm. **G** Representative immunofluorescence images and quantification of BrdU-positive CMs labeled by *Tg(myl7:nucDsRed)* in wild-type (n=11) and *clo^fv087b/+^* (n=10) hearts at 7 dpa. Insets are high magnification of boxed areas. Arrowheads indicate proliferating CMs. Data are the mean ± SEM.; ****p <0.001; unpaired, two-tailed Student’s *t* test. Scale bars, 50 μm. **H** Representative immunofluorescence images and quantification of PCNA-positive CMs and ECs in wild-type (n=10-12) and *clo^fv087b/+^* (n=13-14) hearts at 7 dpa. Arrowheads indicate proliferating CMs or ECs. Data are the mean ± SEM.; ****p <0.001; unpaired, two-tailed Student’s *t* test. Scale bars, 50 μm. **I** Representative immunofluorescence images and quantification of BrdU-positive CMs and ECs in wild-type (n=13) and *clo^m39/+^*(n=11) hearts at 7 dpa. Arrowheads indicate proliferating CMs or ECs. Data are the mean ± SEM.; ***p <0.005, ****p <0.001; unpaired, two-tailed Student’s *t* test. Scale bars, 50 μm. **J** Representative immunofluorescence images of *gata4:eGFP* co-stained with MHC in wild-type (n=14) and *clo^fv087b/+^* (n=15) hearts at 7 dpa. Scale bars, 50 μm.

We next determined whether *naps4l* haploinsufficiency affects the CM proliferation index. By using *Tg(myl7: nucDsRed)* transgenic hearts, we labeled proliferative cardiomyocytes with BrdU molecules. We found the percentage of DsRed^+^/BrdU^+^ proliferative CMs decreased in *clo^fv087b/+^* hearts compared with wild-type siblings at 7 dpa (Figures 1D and 1G). In addition, we applied Mef2 and PCNA antibodies immunostaining to detect CM nuclei with on-going DNA replication, and found that the percentages of Mef2^+^/PCNA^+^ proliferating CMs decreased in *clo^fv087b/+^* hearts at 7 dpa (Figures 1D and 1H). Consistent with this, the percentages of Mef2^+^/BrdU^+^ labeled proliferative CMs also decreased in *clo^m39/+^* mutant hearts (Figures 1D and 1I). Since cardiomyocyte dedifferentiation is another important indicator for myocardial regeneration capacity, we assessed cardiomyocyte dedifferentiation with *Tg(gata4: eGFP)* transgenic reporter (Figure 1D), and found that the GFP expression was significantly reduced in *clo^fv087b/+^*hearts at 7 dpa (Figure 1J). These above data demonstrated that *npas4l* haploinsufficiency caused a defective CM regeneration response after heart injury.

*Cloche* is recognized as the earliest factor for the development of endothelial and hematopoietic cell lineages during zebrafish embryogenesis (Stainier et al., 1995), so we next asked whether *npas4l* haploinsufficiency affects cardiac angiogenesis and endocardial regeneration after heart injury (Figure 1D). Immunostaining data showed that Fli1^+^ ECs in the wound region decreased in both *clo^fv087b/+^* and *clo^m39/+^* mutant hearts (Figures S1B and S1E). This is consistent with the EC proliferation index that the percentages of PCNA^+^/Fli1^+^ ECs, including the endocardium, were reduced in *clo^fv087b/+^* mutant hearts at 7 dpa (Figure 1H); and the percentages of BrdU^+^/Fli1^+^ ECs decreased in *clo^m39/+^* mutant hearts at 7 dpa (Figure 1I). In addition to myocardial and endothelial/endocardial phenotypes in heterozygous *cloche* mutants, we found that the numbers of *coro1a-GFP^+^* leukocytes recruited to the wound region decreased in both *clo^fv087b/+^* and *clo^m39/+^*mutant hearts at 3 dpa (Figures S1B and S1E). All together, these data suggest that in addition to its function in zebrafish embryogenesis (Stainier et al., 1995), *cloche* also plays an important role in adult heart regeneration by modulating CM and EC proliferation and leukocyte recruitment after injury.

### scRNA-seq reveals a potential role of *npas4l* in platelets within the context of cardiac regeneration

To address the cellular and molecular mechanisms regulated by Npas4l, we applied 10x Genomics technology to construct a single-cell transcriptomic atlas of wild-type sibling and *clo^fv087b/+^* mutant hearts in sham-operated zebrafish, and at 3 dpa and 7 dpa. To maintain the cells in the indigenous state and reduce artificial effects caused by cell disassociation, we applied a psychrophilic protease (Adam et al., 2017) and maintained the entire cell disassociation process at 4°C before processing to single-cell sorting and library preparation (Figure 2A). After sequencing and data processing, we clustered and visualized >60,000 single cells in t-SNE dimensionality reduction plots, which consisted of populations of CMs, ECs, epicardium, fibroblasts, immunocytes, erythrocytes, and platelets (Figures 2B, 2C, S2A, S2B and Table S1). Importantly, we found a high enrichment of CMs, which accounted for 44.08% of total cells in our single-cell data. A previous scRNA-seq study had reported that captured CMs displayed much lower transcript counts than non-myocytes, indicating that they might detect transcripts from nuclei rather than cells (Hu et al., 2022). For that reason, we checked in our data and found that the transcript counts and gene counts for CMs were comparable to those of non-myocytes, indicating that we captured intact cells rather than nuclei (Figures S2C and S2D). Among other cell types, we noted two groups of epicardium distinguished by expression of the epicardial activation marker *tcf21* (Figures 2C and S2B). We found an induction of *tcf21^+^* epicardium that was consistent with epicardial activation after injury (Figure S2E). As for immunocyte populations, we acquired T cells, B cells, macrophages, neutrophils, and glial cells. We detected a significant induction of macrophages and neutrophils at 3 dpa, reflecting the inflammatory response to heart injury at early regenerative stages (Figure S2E). Collectively, we constructed a comprehensive single-cell atlas covering the major cell types of regenerating adult zebrafish hearts.

**Figure 2.**
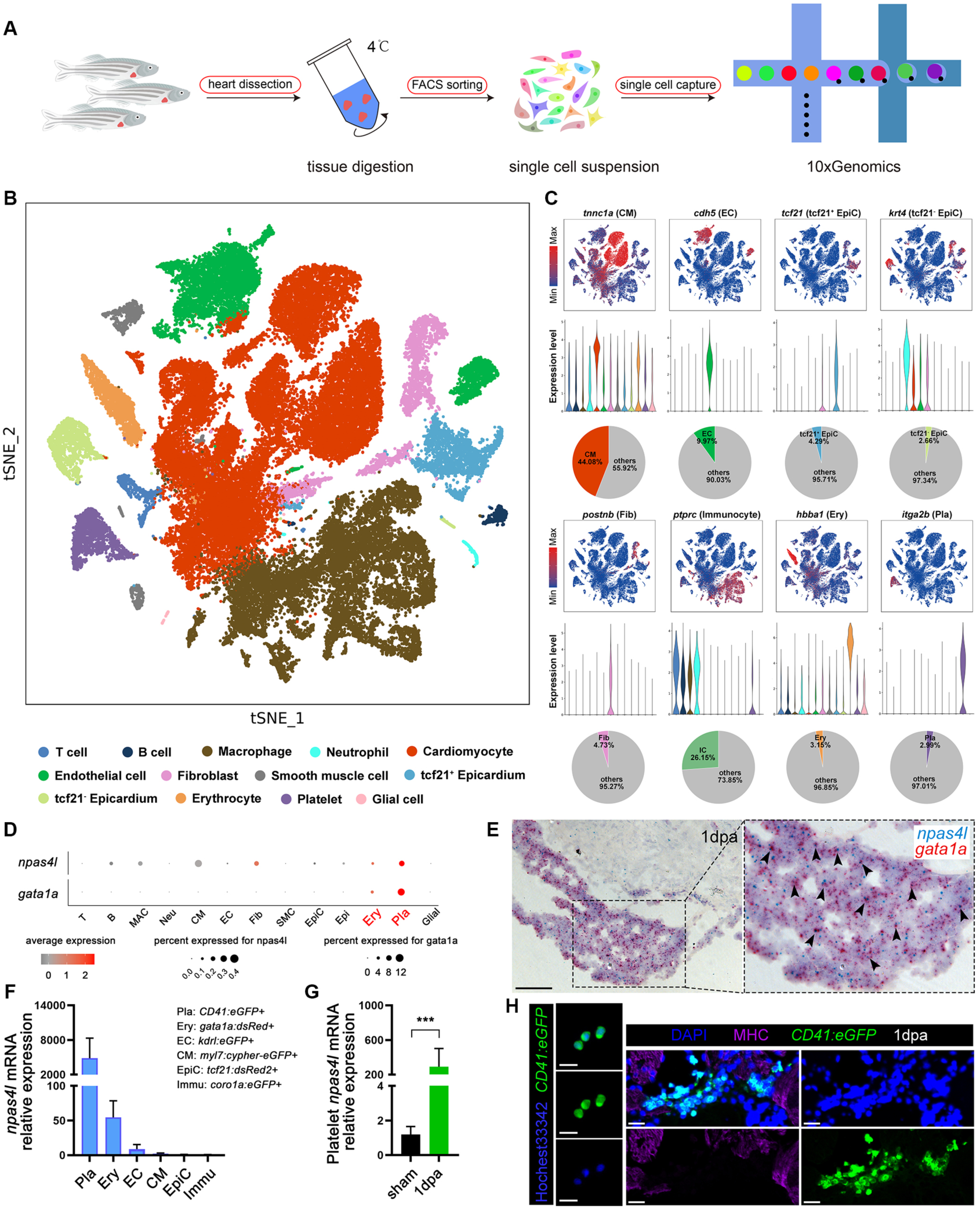
Single-cell RNA-sequencing reveals *npas4l* expression in platelets during heart regeneration. **A** Flowchart of single-cell RNA-sequencing (scRNA-seq) by 10x Genomics of adult zebrafish hearts. **B** t-SNE visualization of clusters of 62,522 cells from both wild-type sibling and *clo^fv087b/+^* mutant hearts. **C** Marker gene expression in each cluster presented in t-SNE (upper) and violin plots (middle), and percentage of each cell type in pie plots (lower). **D** Dot plot showing that *npas4l* and *gata1a* are enriched in erythrocytes and platelets. **E** Representative images of RNAscope *in situ* hybridization with co-staining of *npas4l* and *gata1a* probes in adult zebrafish hearts at 1 dpa. Insets are high magnification of boxed areas. Arrowheads indicate *npas4l^+^* and *gata1a^+^* overlapped signals. Scale bars, 100 μm. **F** Quantitative RT-PCR of *npas4l* mRNA expression in FACS-sorted platelets (*CD41:eGFP*), erythrocytes (*gata1a:dsRed*), endothelial cells (*kdrl:eGFP*), cardiomyocytes (*myl7:cypher-eGFP*), epicardium (*tcf21:dsRed2*), and immunocytes (*coro1a:eGFP*) at 1dpa. mRNA loading was normalized by *rpl13a*. n=3 replicates per group, and around 6000 cells pooled together pre replicate. Data are the mean ± SEM. **G** Quantitative RT-PCR of *npas4l* mRNA expression in FACS-sorted platelets (*CD41:eGFP*) at sham and 1dpa. mRNA loading was normalized by *rpl13a*. n=3 replicates per group, and around 6000 cells pooled together per replicate. Data are the mean ± SEM.; ***p<0.005; unpaired, two-tailed Student’s t test. **H** Left: Representative fluorescence images of isolated platelets (*CD41:eGFP*) stained with Hochest33342. Scale bars, 10 μm. Right: Representative immunofluorescence images of *CD41:eGFP* co-stained with MHC and DAPI on zebrafish heart sections at 1 dpa. Scale bars, 20 μm.

To determine the *npas4l-*expressing cell type, we sought *npas4l-*expressing cells in our single-cell data and found 207 cells that had *npas4l* transcripts. By plotting the cell ratio and expression level of *npas4l*, we observed a significant enrichment of *npas4l* expression in both platelets and erythrocytes, which are the same with *gata1a*-expressing cell type (Figure 2D). Although we noted weak *npas4l* expression in other cell types like fibroblasts and CMs, the expression levels or percentages were not comparable with those in platelets. To confirm the *npas4l-*expressing cell types, we also designed a RNAscope probe targeted against *npas4l* and applied it to whole-mount RNAscope *in situ* hybridization of 2-somite stage zebrafish embryos. The visualized signals mirrored the *npas4l* expression pattern reported previously (Reischauer et al., 2016) (Figure S3A). By using this probe to perform RNAscope *in situ* hybridization on adult heart sections after injury, we found that the signal intensity of *npas4l* was nearly reduced by half in *clo^fv087b/+^* hearts compared with wild-type sibling hearts at 1 dpa (Figure S3B). We further assessed the dynamic changes of *npas4l* upon heart injury, we found that a large amount of *npas4l* signals were detected in the infarct area at 3 dpa and the signals started to decrease subsequently (Figure S3C). We also checked *npas4l* expression window by performing quantitative RT-PCR of the whole ventricle after heart amputation. We found that *npas4l* showed an early response to injury, evident within 1 hour post amputation (1 hpa), peaking at 1 dpa, and subsequently declining thereafter (Figure S3D). All of these data suggested *npas4l* acted as an early player during heart regeneration, which temporally matched injury-triggered onset of thrombosis and its subsequent degradation. Then, by utilizing cell-type specific transgenic reporter lines, including *Tg(CD41:eGFP)*, *Tg(gata1a: dsRed)*, *Tg(kdrl:eGFP)*, *Tg(myl7: cypher-eGFP)*, *Tg(tcf21: dsRed2)* and *Tg(coro1a:eGFP)*, We harvested, dissociated, and FACS sorted cells from hearts at 1dpa, including platelets, erythrocytes, endothelial cells, cardiomyocytes, epicardium, and immunocytes. RT-PCR analysis revealed a remarkable enrichment of *npas4l* expression in platelets, followed by erythrocytes, while hardly in endothelial cells, cardiomyocytes, epicardium, and immunocytes (Figure 2F). By using RNAscope *in situ* hybridization on adult heart sections after injury, we found that, consistent with our single-cell data, *npas4l* signals shared a remarkable co-localization with *gata1a*, which labeled erythrocytes and platelets and were densely present within the blood clot area after injury (Figure 2E). Notably, at the wound site, *npas4l* was enriched in *itga2b^+^* platelets in close proximity to the edge of the border zone (Figure S4A). Furthermore, we found no evident co-localizations of *npas4l* with other heart cell markers, including *postnb* (fibroblasts), MHC (CMs), *kdrl* (ECs), *coro1a* (immunocytes), and *tcf21* (epicardium) (Figures S4B-S4D), suggesting that *npas4l* expression was highly enriched in platelets and erythrocytes. When we compared *npas4l* expression in platelets before and after heart injury, we found dramatic induction of *npas4l* expression in platelets after heart injury (Figure 2G). Unlike mammals, teleost fishes including zebrafish preserved nucleated platelets and erythrocytes evolutionarily which made transcriptional regulation feasible (Bronnimann et al., 2018; Liu et al., 2007). And we confirmed this platelet nucleation in zebrafish by co-localization of *Tg(CD41:eGFP)* platelets with nuclear indicators DAPI and Hochest33342 by immunostaining and fluorescent imaging (Figure 2H). Taken together, we established a single-cell atlas of regenerating adult zebrafish hearts and identified *npas4l* as an early regenerative factor derived from *gata1a^+^* blood cells, particularly platelets at the wound site.

### Platelets are essential for zebrafish heart regeneration

To further address the critical role of platelets in zebrafish heart regeneration, we applied *Tg(CD41:eGFP)* transgenic reporter to label platelets, and found that platelets coagulated in the wound region neighboring the myocardium right after injury and diminished from 1 to 7 dpa (Figure 3A). Because amputation is associated with profuse bleeding, we investigated the functions of platelets beyond coagulation by examining regeneration following NTR-mediated genetic ablation of CMs (Curado et al., 2007). Using *Tg(myl7:ECFP-NTR)*, in which NTR was overexpressed specifically in CMs, we first confirmed CMs ablation efficiency in this model by immunostaining (Figure S5A). Our CM ablation protocol revealed CMs were sufficiently damaged, showing weaker signal intensities of myosin heavy chain (CM cytoplasm marker) and myocyte-specific enhancer factor 2 (CM nuclei marker) along with a disrupted sarcomere structure labeled by α-Actinin after metronidazole treatment (Figure S5C). Importantly, we found that *itga2b^+^* platelets accumulated in the heart beginning at 3 days after induction of NTR with metronidazole treatment (dpt), peaked at 7 dpt, and returned to baseline levels by 14 dpt (Figure 3B). These injury-induced responses and recruitment of platelets implicated a potential regenerative role of platelets beyond their function in blood clotting. By using NTR-mediated CM ablation model, we also found that both Mef2^+^/BrdU^+^ CMs and Fli1^+^/BrdU^+^ ECs decreased in *clo^fv087b/+^*mutant hearts at 7 dpt (Figure S5B and S5D), which is consistent with the data obtained from ventricular amputation model in zebrafish.

**Figure 3.**
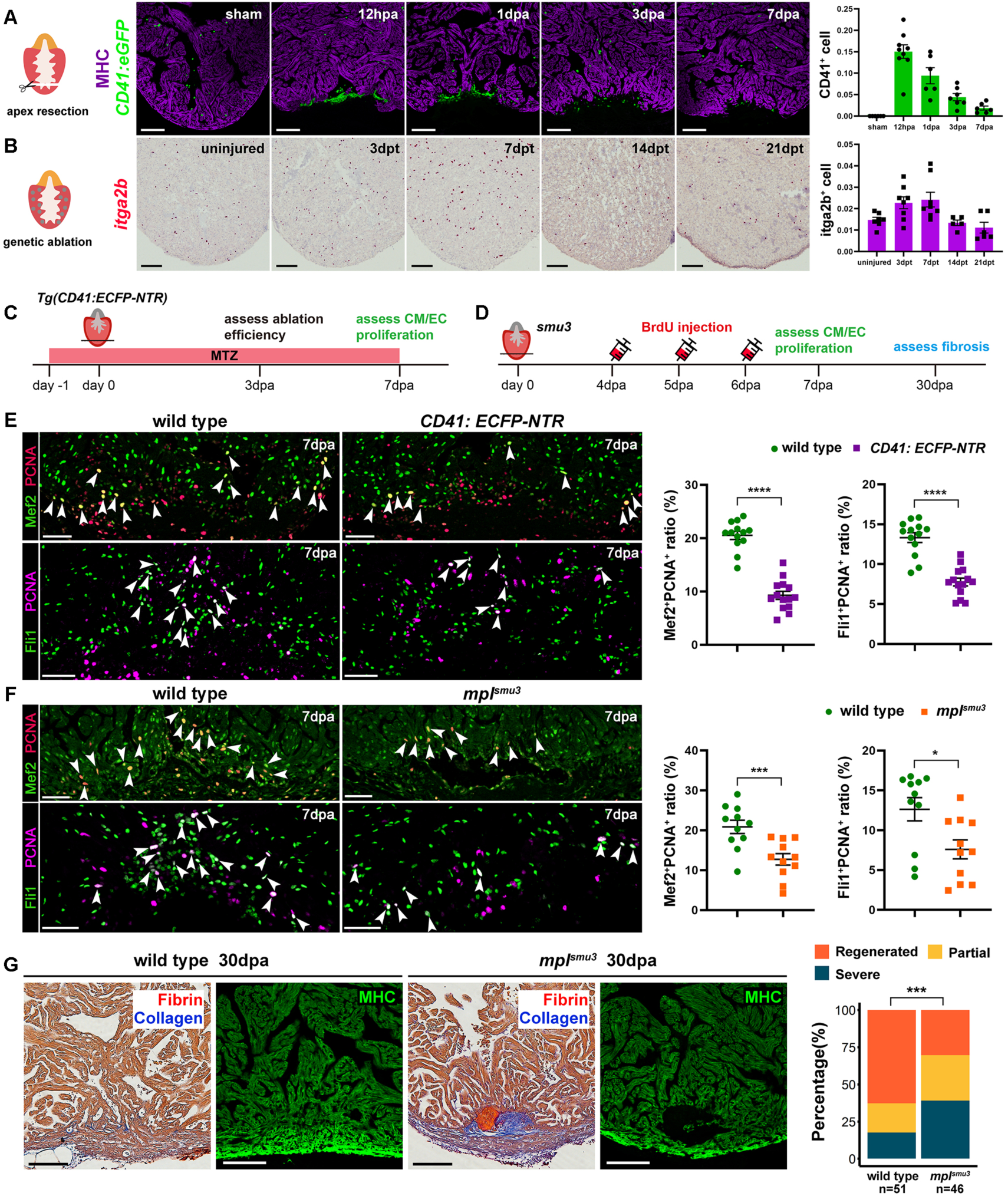
Injury-induced platelets are essential for heart regeneration. **A** Left, schematic of apex resection. Right, representative immunofluorescence images and quantification of *CD41:eGFP* (MHC immunofluorescence signal labels the myocardium) in zebrafish hearts from sham, and at 12 h, 1 day, 3 days, and 7 days post amputation. n=6-9 hearts per group. Data are the mean ± SEM. Scale bars, 100 μm. **B** Left, schematic of genetic ablation of CMs using *Tg(myl7:ECFP-NTR)* transgenic zebrafish. Right, representative images of RNAscope *in situ* hybridization and quantification of *itga2b^+^* platelets in *myl7:ECFP-NTR* transgenic hearts in uninjured, and at 3 to 21 days post MTZ treatment (dpt). n=5-8 hearts per group. Data are the mean ± SEM. Scale bars, 100 μm. **C, D** Schematic diagram of experimental design to explore platelets function in *Tg(CD41:ECFP-NTR)* (**C**) and *mpl^smu3^* (**D**). **E** Representative immunofluorescence images and quantification of PCNA-positive CMs and ECs in wild-type sibling (n=13) and *CD41:ECFP-NTR* (n=14) transgenic hearts at 7 dpa. Arrowheads indicate proliferating CMs or ECs. Data are the mean ± SEM.; ****p <0.001; unpaired, two-tailed Student’s *t* test. Scale bars, 50 μm. **F** Representative immunofluorescence images and quantification of PCNA-positive CMs and ECs in wild-type sibling (n=11) and *mpl^smu3^* (n=11) mutant hearts at 7 dpa. Arrowheads indicate proliferating CMs or ECs. Data are the mean ± SEM.; *p <0.05; ***p <0.005; unpaired, two-tailed Student’s *t* test. Scale bars, 50 μm. **G** Representative images and quantification of AFOG staining and MHC immunofluorescence of wild-type (n=51) and *mpl^smu3^* (n=46) heart sections at 30 dpa. Data represents the percentage of each group; ***p <0.005; Pearson chi-square test. Scale bars, 100 μm.

To directly address platelet function during heart regeneration, we investigated heart regeneration processes by depleting platelets by either NTR-mediated ablation of platelets or platelet-deficient mutant *mpl^smu3^*(Lin et al., 2017). Considering the fact that *gata1a^+^* blood cells consisted mostly of platelets in our single-cell data (Figure 2D), we generated *Tg(gata1a:ECFP-NTR)* zebrafish in which NTR was overexpressed in *gata1a^+^* blood cells. By using RNAscope *in situ* hybridization, we found that about half of *itga2b^+^* platelets were eliminated in *Tg(gata1a:ECFP-NTR)* hearts at 3 dpt (Figure S6A and S6B), and a notable decrease of both PCNA^+^/Mef2C^+^ CMs and PCNA^+^/Fli1^+^ ECs in *Tg(gata1a:ECFP-NTR)* hearts at 7 dpa (Figure S6A and S6C). To achieve a more specific depletion of platelets, we also generated *Tg(CD41:ECFP-NTR)* zebrafish, in which NTR was overexpressed in *CD41^+^* platelets. By using RNAscope *in situ* hybridization to validate platelet ablation efficiency, we found that nearly all *itga2b^+^* platelets were eliminated in *Tg(CD41:ECFP-NTR)* hearts at 3 dpa (Figures 3C and S7A), and consistent with the phenotype observed in *Tg(gata1a:ECFP-NTR)* hearts, we found a notable decrease of both PCNA^+^/Mef2^+^ CMs and PCNA^+^/Fli1^+^ ECs in *Tg(CD41:ECFP-NTR)* hearts at 7 dpa (Figures 3C and 3E). Additionally, by using a platelet-deficient *mpl^smu3^* mutant, in which >90% of platelets were depleted in early embryos (Lin et al., 2017), we also found few *itga2b^+^* platelets in the infarct area of mutant hearts after ventricular resection (Figure S7B). Consistent with the NTR-based deletion of platelets above, we found that both BrdU and PCNA labeled proliferative CMs and ECs decreased in *mpl^smu3^* mutant hearts at 7 dpa (Figures 3D, 3F and S7C). Moreover, AFOG and MHC immunofluorescence staining data showed larger scars consisting of fibrin and collagen and less myocardial regeneration in mutant hearts compared with wild-type sibling hearts at 30 dpa (Figures 3G).

To further assess whether *npas4l* influenced the formation of platelets, we quantified and found no difference on the numbers of *Tg(CD41:eGFP)*-labeled platelets between *clo^fv087b/+^* and wild-type hearts at 1dpa (Figure S8A). Furthermore, we overexpressed *npas4l* (Npas4l OE) in *gata1a*-positive cells by utilizing the *Tg(ubi:loxp-eGFP-stop-loxp-npas4l)* [designated *Tg(ubi-npas4l)*] and *Tg(gata1a:Cre-ERT2)* double transgenic fish lines. By utilizing RNAscope *in situ* hybridization with *cre* and *gata1a* probes, we confirmed *cre* was specifically expressed in *gata1a^+^* erythrocytes after heart injury (Figure S8B). Following 4-hydroxytamoxifen (4-HT) treatment, we extracted and isolated total RNA of *gata1a^+^* platelets/erythrocytes or the whole ventricles from *Tg(ubi-Npas4l)* control and Npas4l OE hearts at 3- or 7-days post 4-HT treatment. Quantitative RT-PCR confirmed that the levels of *npas4l* mRNA increased in both ventricles and isolated platelets/erythrocytes in Npas4l OE hearts, indicating *npas4l* was successfully overexpressed in platelets/erythrocytes (Figure S8C). By using RNAscope *in situ* hybridization with *itga2b* probes, we found overexpression of *npas4l* had no evident effect on the numbers of *itga2b^+^*platelets in Npas4l OE hearts compared to *Tg(ubi-npas4l)* control hearts at 7 dpt (Figure S8D). Together, these data supported that Npas4l is essential for injury-induced platelet activation and function during zebrafish heart regeneration, but is not critical for formation and recruitment of platelets in the injured hearts.

### Npas4l regulates a platelet ligand network program to modulate cell-cell interactions

To address how Npas4l in platelets regulates heart regeneration, we revisited our scRNA-seq datasets and focused on ligand-receptor activity among different heart cell types. By plotting the ligand expression of each cluster identified in zebrafish hearts, we found that more ligands were expressed in platelets, smooth muscle cells, and macrophages, suggesting that these cells were actively secretory (Figure 4A). We noted several genes reported to be connected with heart regeneration as expressed in platelets, including *vegfc* (El-Sammak et al., 2022), *Serpine1* (Munch et al., 2017), and *bmp6*. Bmp6 has been described to function downstream of Tbx20, mediating EC proliferation (Fang et al., 2020). By comparing the numbers of ligand-receptor pairs in each cluster between wild-type and *clo^fv087b/+^* mutant hearts during regeneration, we found an evident decrease in ligand-receptor pairs in *clo^fv087b/+^*hearts, indicating the weaker cell-cell interactions and signal transduction in the mutant hearts (Figure 4B). As for platelet interactions with other cells, since we did not detect any significant quantitative changes of ligand-receptor pairs between wild-type siblings and *clo^fv087b/+^*mutant hearts, we switched our focus to the ligand-receptor activities in platelets that mediated interactions with CMs and ECs. By sub-clustering platelets, CMs, and ECs, we obtained 6 subsets of platelets, 11 subsets of CMs, and 7 subsets of ECs (Figures S9-S11 and Tables S2-S4). In addition, we found dynamic changes in these subsets in *clo^fv087b/+^* hearts at 3 dpa and 7 dpa (Figures S9C, S10C and S11C). Importantly, by plotting the ligand-receptor expression of each subcluster, we noted 3 highly expressed ligand-receptor pairs in platelet-CM interactions including HBEGF-EGFR, BMP6-BMPR1A-ACTR2A, and BMP6-BMPR1A-BMPR2, which mainly targeted CM3 and CM9 by all 6 platelet subsets (Figure 4C). And we measured a decrease of the CM3 and CM9 ratio in *clo^fv087b/+^* hearts at 3 dpa and 7 dpa (Figure S9C). By analyzing their characteristics and cellular functions, we found that CM3 (marked by *myh10*) and CM9 (marked by *myh6*) were enriched with pathways related to cardiac muscle cell differentiation and proliferation, indicating that these subsets represent potential proliferative CMs (Figures S9D-S9F and Table S2). Notably, “Transforming growth factor beta1 production” and “Regulation of Smad protein complex assembly” were also included in CM9, suggesting that CM9 is enriched in Bmp6-mediated TGF-beta signals. In line with platelet-CM interactions, we noted 3 highly expressed ligand-receptor pairs in platelet-EC interactions including VEGFC-KDR, VEGFC-FLT4. and HBEGF-EGFR, which mainly targeted EC2 and EC7 (Figure 4D). Consistent with this, we observed a decrease of both EC2 and EC7 ratio in *clo^fv087b/+^* hearts at 3 dpa and 7 dpa (Figure S10C). Analysis of the characteristics and cellular functions of both groups demonstrated that EC2 (marked by *plvapb*) and EC7 (marked by *cxcl12a*) were enriched in vascular EC proliferation or migration (Figures S10D-S10F and Table S3). Furthermore, we noted pathways enriched in EC7 like “Lympho-angiogenesis” and “Regulation of retinoic acid receptor signaling pathway”, which have been reported to participate in angiogenesis and heart regeneration. Both clues suggested that EC2 and EC7 were potential proliferative EC subsets. Together, our scRNA-seq revealed platelets, smooth muscle cells, and macrophages as major secretory cell clusters, and Bmp6, Hbegf, and Vegfc as key platelet ligands in mediating platelet interactions with CMs and ECs.

**Figure 4.**
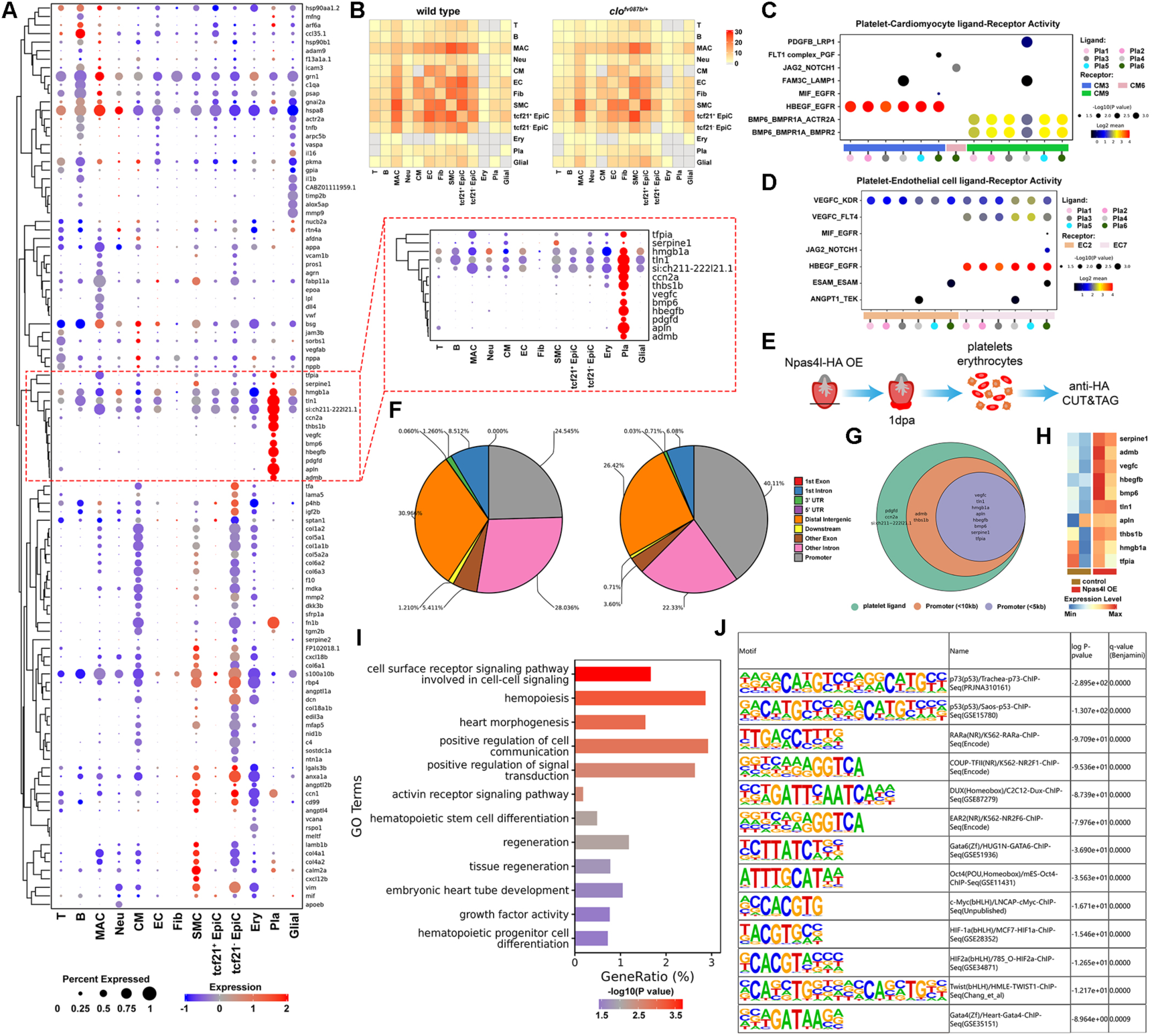
Npas4l regulates a platelet ligand network program to modulate cell-cell interactions. **A** Dot plot showing ligand expression in each cell type of the zebrafish heart. The inset is high magnification of the boxed area highlighting platelet-enriched ligands. **B** Heatmap depicting the numbers of ligand-receptor pairs among cell clusters in wild-type and *clo^fv087b/+^*hearts during regeneration. **C, D** Dot plot showing significant ligand-receptor activity (P <0.05) between platelet-CM (**C**) and platelet-EC (**D**). **E** Flowchart of Npas4l-HA ChIP-seq by CUT&TAG during heart regeneration. **F** Pie plots showing Npas4l protein binding peaks distribution on the genome scale, of which the promoter is defined within ±5kb (left) or ±10 kb (right) to TSS. **G** Venn diagram of platelet-enriched ligands with Npas4l-regulated ligands. **H** Heatmap presenting the expression of Npas4l-regulated ligands in *gata1a^+^*platelets/erythrocytes between Npas4l OE and control hearts at 7 dpt. n=2 replicates per group, and around 6000 cells pooled together per replicate. **I** Gene ontology analysis showing pathways directly regulated by Npas4l. **J** Motif enrichment presenting preference of Npas4l binding sequences.

Since Npas4l is a classic basic helix-loop-helix transcription factor and only one report identified its direct binding targets during zebrafish embryogenesis (Marass et al., 2019), we decided to assess its targets during heart regeneration. By generating a transgenic line overexpressing HA tagged Npas4l (Npas4l-HA OE) consisting of *Tg(ubi:loxp-stop-loxp-Npas4l-HA)* and *Tg(gata1a:Cre-ERT2)*, we harvested 1-dpa hearts, isolated the platelets/erythrocytes, and performed the high-sensitive ChIP-seq with CUT&TAG method using anti-HA antibody (Figure 4E). By comparing Npas4l-HA OE hearts with *Tg(gata1a:Cre-ERT2)* control hearts, we got *npas4l*-specific binding peaks. To calculate their regional proportion on the whole genome scale, we chose two ranges to define promoter regions, a narrow range (±5kb to TSS) and a broad range (±10kb to TSS). The distribution pie plot revealed that promoter, first intron, and distal intergenic regions were dominantly anchored by *npas4l* binding peaks (Figure 4F). Moreover, among all the platelet-specific ligands, *npas4l* showed promoter binding activities on most of these genes, and some of them including *bmp6*, *vegfc*, *hbegfb*, *serpine1*, *tfpia*, *hmgb1a*, *apln*, and *tln1* were included in a narrow range of promoter-regulated gene scale (Figure 4G). To further confirm Npas4l regulation on these genes expression, we isolated *gata1a^+^* platelets/erythrocytes from Npas4l OE and control hearts after 7 days post 4-HT treatment (7dpt) and analyzed the transcriptome by RNA-sequencing. Notably, we found a fully upregulated RNA expression of these ligand genes, indicating that *npas4l* directly bound their promoters and regulated their transcription levels (Figure 4H and Table S7). The gene ontology analysis of *npas4l* binding peaks revealed that, in addition to well-known biological process like “hemopoiesis” and “hematopoietic progenitor cell differentiation”, *npas4l* also regulated many pathways related to cell communication, signal transduction, growth factor activity, and tissue regeneration, which correlated well with the conclusion above that platelet *npas4l* directly regulated ligands expression to mediated cell-cell interactions with CMs and ECs (Figure 4I and Table S5). Furthermore, by motif analysis of Npas4l binding peaks, we noticed a significant HIF1α consensus sequence along with other pluripotent genes recognition motifs, including Oct4, c-Myc and Gata6 (Figure 4J). On top of these motifs, the tumor-related protein p73 and p53 recognition motifs lied up, which suggested *npas4l* might synergistically cooperate with these proteins to regulate their target genes (Figure 4J). But this hypothesis warrants further investigations in the future. Taken together, our data suggest that Npas4l directly regulates a platelet ligand network program to coordinate cell-cell interactions during heart regeneration.

### Platelet Npas4l regulates BMP6 signaling for CM proliferation and regeneration

Since we found that neither loss-of-function nor gain-of-function of *npas4l* affects the platelet numbers and responses to injury (Figure S8A and S8D), we then asked how loss-of-function and gain-of-function of *npas4l* affects platelet transcriptomic profiling. By sorting out *CD41^+^* platelets from wild type siblings and *clo^fv087b/+^* mutants, we performed bulk RNA-sequencing, compared the transcriptomic changes with RNA-seq data of Npas4l OE hearts mentioned above, and identified Npas4l-regulated genes and pathways. We discovered 1626 differentially expressed genes (DEGs) between wild-type siblings and *clo^fv087b/+^* mutant hearts at 1 dpa, and 1874 DEGs between control and Npas4l OE hearts at 7 dpt, of which 269 genes were shared by both cases (Figure 5A and Table S7). By categorizing these genes based on expression changes dependent on *npas4l*, we identified two groups of genes that were assigned as Npas4l-positively or -negatively regulated genes (Figure 5B). Notably, we found that *bmp6* along with other TGF-beta superfamily members like *rock1*, *bambib*, and *tgfb1b* were all included in the Npas4l-positively regulated genes. Analysis of pathways and biological processes also noted that the TGF-beta signaling pathway was significantly enriched in Npas4l-positively regulated pathways (Figures 5C, S12 and Table S7). Moreover, genome browsing of *npas4l* binding peaks and read counts near *bmp6* transcription start site, we observed significant enriched peaks located within the promoter and first intron of two *bmp6* transcripts, indicating of *npas4l* direct binding and regulation (Figure 5E). Since *bmp6* was found to be one of the key platelet ligands mediating crosstalk between platelets and CMs (Figure 4C), these data suggested *bmp6* as a potential downstream effector of Npas4l. We then examined the *bmp6* expression pattern after injury and found that injury triggered *bmp6* elevation from 1 dpa to 7 dpa, and in the infarct area, *bmp6* expression was restricted to *itga2b^+^* platelets only; while in the border and remote zones, we also found *bmp6* expression in ECs (Figures 5D, S13 and S14A), which was consistent with a previous report (Fang et al., 2020). To further validate Npas4l regulation of *bmp6*, we assessed the expression levels of *bmp6* and other platelet ligand mRNAs in platelets between wild-type siblings and *clo^fv087b/+^* mutant hearts at 1 dpa. We found an evident reduction of *bmp6*, *tfpia*, *thbs1b*, and *vegfc* mRNA expression and a slight increase of *apln* mRNA expression in *clo^fv087b/+^* hearts (Figure 5F). In platelets/erythrocytes from Npas4l OE hearts at 7 dpt, we found a consistent increased expression of these platelet ligand mRNAs in which *bmp6* expression had the most striking induction (Figure 5G), emphasizing the Npas4l regulation of *bmp6*. In addition, we discovered that Bmp6 protein expression was reduced in *clo^fv087b/+^* hearts at 3 dpa by western blots (Figure 5H). RNAscope *in situ* hybridization of *bmp6* revealed a remarkable increase of the *bmp6* signal in Npas4l OE hearts compared with control hearts at 7 dpt (Figure 5I). All these data together supported the conclusion that platelet *bmp6* was positively regulated by Npas4l.

**Figure 5.**
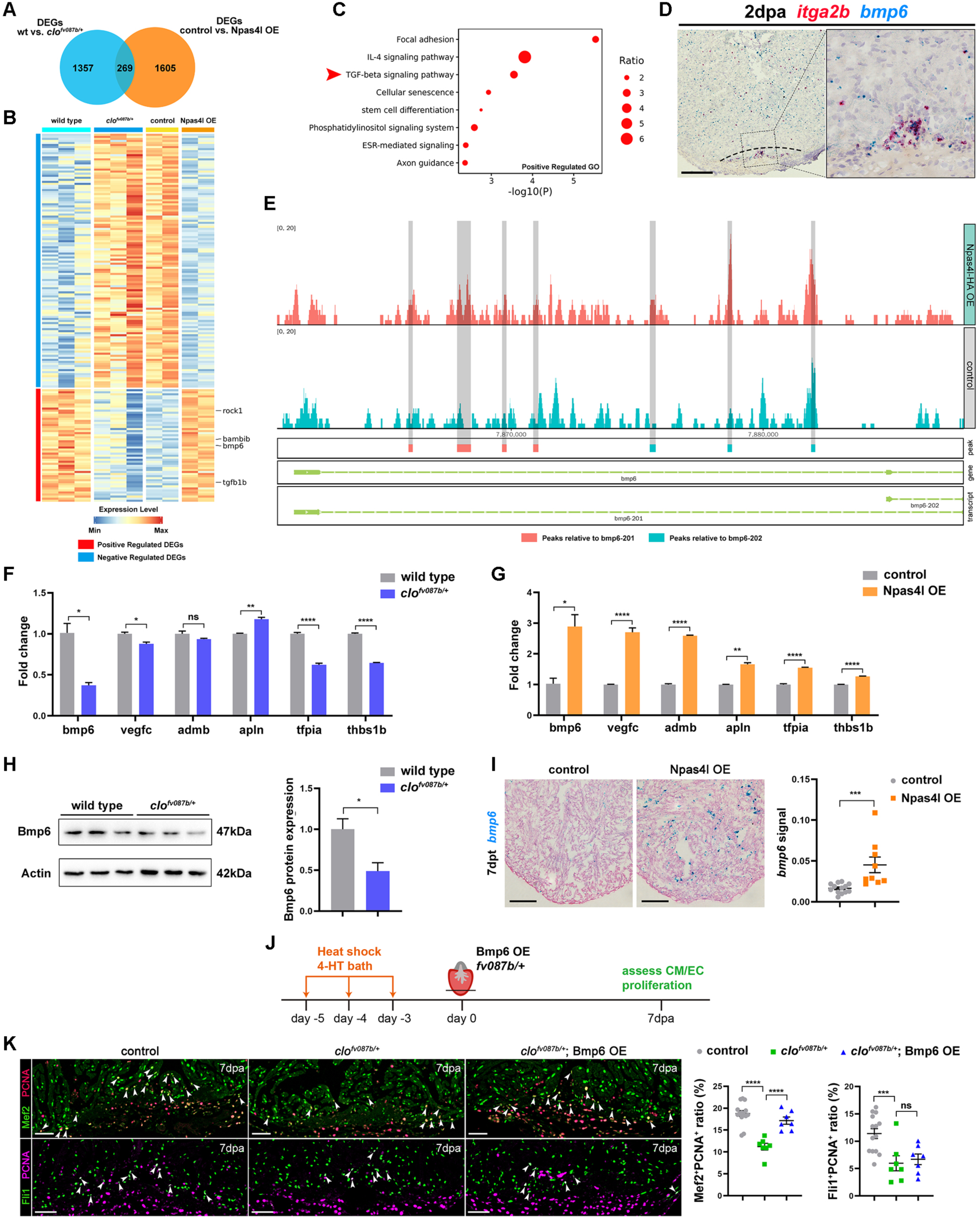
Platelet Npas4l regulates BMP6 signaling for CM proliferation and regeneration. **A** Venn diagram of bulk RNA-seq data showing the overlap of differentially expressed genes (DEGs) from *CD41^+^* cells (wild-type sibling *vs clo^fv087b/+^* mutant hearts at 1 dpa, n=3 replicates per group, around 6000 cells pooled together per replicate) and *gata1a^+^*cells (control sibling *vs* Npas4l OE hearts at 7 dpt, n=2 replicates per group, around 6000 cells pooled together per replicate). Fold change >1.4; p-value adjusted (p_adj_) <0.05. **B** Heatmap showing up-regulated genes (red) and down-regulated genes (blue) by Npas4l from the overlapped DEGs in panel A. **C** Pathway analysis showing enriched terms of Npas4l positively-regulated genes. The arrowhead indicates the BMP6-related TGF-beta signaling pathway. **D** Representative images of RNAscope *in situ* hybridization using *bmp6* probe co-stained with *itga2b* probe in zebrafish hearts at 2 dpa. The inset is high magnification of the boxed area. Dashed lines delineate the wound edge. Scale bar, 100 μm. **E** Browser tracks proximal to *bmp6* transcription start site, indicating anti-HA CUT&TAG enriched read counts and Npas4l binding peaks of isolated platelets/erythrocytes from Npas4l-HA OE and control hearts at 1 dpa. **F, G** Quantitative RT-PCR of platelet ligands mRNA expression normalized by *rpl13a* in CD41-positive cells (wild type *vs clo^fv087b/+^*hearts at 1 dpa, **F**) and gata1a-positive cells (control *vs* Npas4l OE hearts at 7 dpt, **G**) from zebrafish hearts. One of three independent experiments is presented, n=3 replicates per group, around 6000 cells pooled together per replicate. Data are the mean ± SEM.; *p <0.05; *p <0.01; ****p <0.001; unpaired, two-tailed Student’s *t* test. **H** Western blot and quantification of Bmp6 expression in wild-type and *clo^fv087b/+^*hearts at 1 dpa. n=3 replicates per group, 10 hearts pooled together per replicate. β- Actin serves as loading control. Data are the mean ± SEM.; *p <0.05; unpaired, two-tailed Student’s *t* test. **I** Representative images of RNAscope *in situ* hybridization and quantification of *bmp6* in control (n=13) and Npas4l OE (n=9) hearts at 7 dpt. Data are the mean ± SEM.; ***p <0.005; unpaired, two-tailed Student’s *t* test. Scale bars, 100 μm. **J** Schematic diagram of experimental design to explore whether *bmp6* over-expression rescues *clo^fv087b/+^* phenotype. **K** Representative immunofluorescence images of PCNA-positive CMs and ECs in *hsp70:Cre-ERT2* (control, n=14), *hsp70:Cre-ERT2; clo^fv087b/+^* (*clo^fv087b/+^*, n=7), and *clo^fv087b/+^; hsp70:Cre-ERT2; ubi:loxp-stop-loxp-bmp6* (*clo^fv087b/+^*, Bmp6 OE) (n=7) hearts at 7 dpa. Arrowheads indicate proliferating CMs or ECs. Data are the mean ± SEM.; ***p <0.005; ****p <0.001; one-way ANOVA with LSD test. Scale bars, 50 μm.

To further explore Bmp6 function in heart regeneration, we applied CRISPR/Cas9 genome editing technology to generate a *bmp6* mutant line, of which 1.6kb sequence between exon3 and exon4 was deleted, resulting in a frame shift and pre-terminated protein translation (Figure S14B). Western blot and quantitative RT-PCR results revealed that both mRNA and protein expression of *bmp6* was significantly downregulated in *bmp6* mutant, while other major BMP ligands mRNA including *bmp2a*, *bmp2b*, *bmp7a* and *bmp7b* remained unaffected (Figure S14C-S14E). Importantly, by comparing with wild type hearts, *bmp6* mutant hearts exhibited a remarkable decrease of BrdU-positive CMs but not ECs at 7 dpa (Figure S14F). Besides, we generated a *bmp6* over-expression line (Bmp6 OE) consisting of *Tg(ubi:loxp-stop-loxp-bmp6)* and *Tg(hsp70:Cre-ERT2)* and asked whether *bmp6* over-expression rescued *clo^fv087b/+^* mutant phenotype. Following heat shock and 4-HT treatment, we observed *bmp6* over-expression significantly rescued PCNA-positive CMs index in *clo^fv087b/+^* mutant hearts at 7 dpa but had no effects on PCNA-positive ECs (Figure 5J and K). Altogether, these data suggested that Bmp6 acts downstream to Npas4l in platelets to regulate cardiomyocyte proliferation.

### Platelet Npas4l is both necessary and sufficient for heart regeneration

Since we identified *npas4l* as a platelet-derived regulator of cardiomyocyte proliferation, we asked whether over-expression of *npas4l* in platelets was sufficient to promote quiescent CMs to enter the cell cycle. The immunofluorescence on uninjured heart sections showed that over-expression of *npas4l* in *gata1a^+^* platelets caused significant induction of proliferative CMs and ECs in Npas4l OE hearts at 7 days after 4-HT treatment (dpt) (Figures S15A and S15D). By crossing the Npas4l OE line with *clo^fv087b/+^*, we found that the percentages of BrdU^+^ proliferative CMs and ECs decreased in *clo^fv087b/+^* hearts compared with wild-type control hearts, and this was rescued by overexpression of *npas4l* in *clo^fv087b/+^; Npas4l OE* hearts at 7 dpa (Figures S15B and S15E). Furthermore, we generated a platelet-specific *npas4l* over-expression line (Pla Npas4l OE) consisting of *Tg(ubi:loxp-stop-loxp-Npas4l-HA)* and *Tg(CD41:Cre-ERT2)*. RNAscope *in situ* hybridization with *cre* and *itga2b* probes showed that *cre* was specifically expressed in *itga2b^+^* platelets after injury (Figure S16). By crossing the Pla Npas4l OE line with *clo^fv087b/+^*, we asked whether platelet-specific *npas4l* over-expression rescued CM proliferation defects in *clo^fv087b/+^* mutant, or even reinforced the CM proliferation intensity in uninjured wild-type hearts.

Following the 4-HT bathing and apex amputation, we found an evident restore of CM and EC proliferation index nearly to the control level in *clo^fv087b/+^* hearts after *npas4l* over-expression in platelets at 7 dpa (Figures 6A-6C). More importantly, over-expression of *npas4l* in platelets strengthened the proliferation capacities of CMs and ECs after heart injury compared with the control hearts, indicating platelet *npas4l* was potent to trigger the cycling in zebrafish heart regeneration (Figures 6A-6C). Taking a step further, we applied Pla Npas4l OE consisting of *Tg(ubi:loxp-GFP-stop-loxp-Npas4l)* and *Tg(CD41:Cre-ERT2)* to ask whether over-expression of *npas4l* in platelets was sufficient to promote quiescent CMs and ECs to enter the cell-cycle. Strikingly, over-expression of *npas4l* in platelets showed significant induction of proliferative CMs in both compact layer and trabecular layer at 7 days post 4-HT treatment (dpt) (Figures 6A and 6D). On the outer layer of the ventricle, we also found increased cycling activities of coronary endothelial cells in Pla Npas4l OE hearts at 7 dpt, indicating over-expression of *npas4l* in platelets also promoted angiogenesis in zebrafish hearts (Figures 6A and 6D). Since *bmp6* functioned downstream of *npas4l*, we asked whether Bmp6 signaling played a role in promoting quiescent CMs and ECs into the cell-cycle. By injecting the BMP signaling inhibitors K02288 or LDN193189 into control and Npas4l OE hearts, we found that inhibition of BMP signaling partially ablated PCNA^+^ CMs in Npas4l OE hearts (Figure S15F). As for ECs, we found that BMP inhibitors caused a slight reduction in EC proliferation index but this failed to reach statistical significance. In addition, by genetically overexpressing *bmp6* in zebrafish hearts, we found no effects on promoting quiescent CM and EC cycling (Figure S17), indicating that although *bmp6* contributed to *npas4l*-induced cardiomyocyte cycling, other Npas4l-regulated factors are required for inducing cardiomyocyte proliferation and regeneration. Together, these results suggested that platelet *npas4l* is required and sufficient for promoting CM and EC proliferation and heart regeneration.

**Figure 6.**
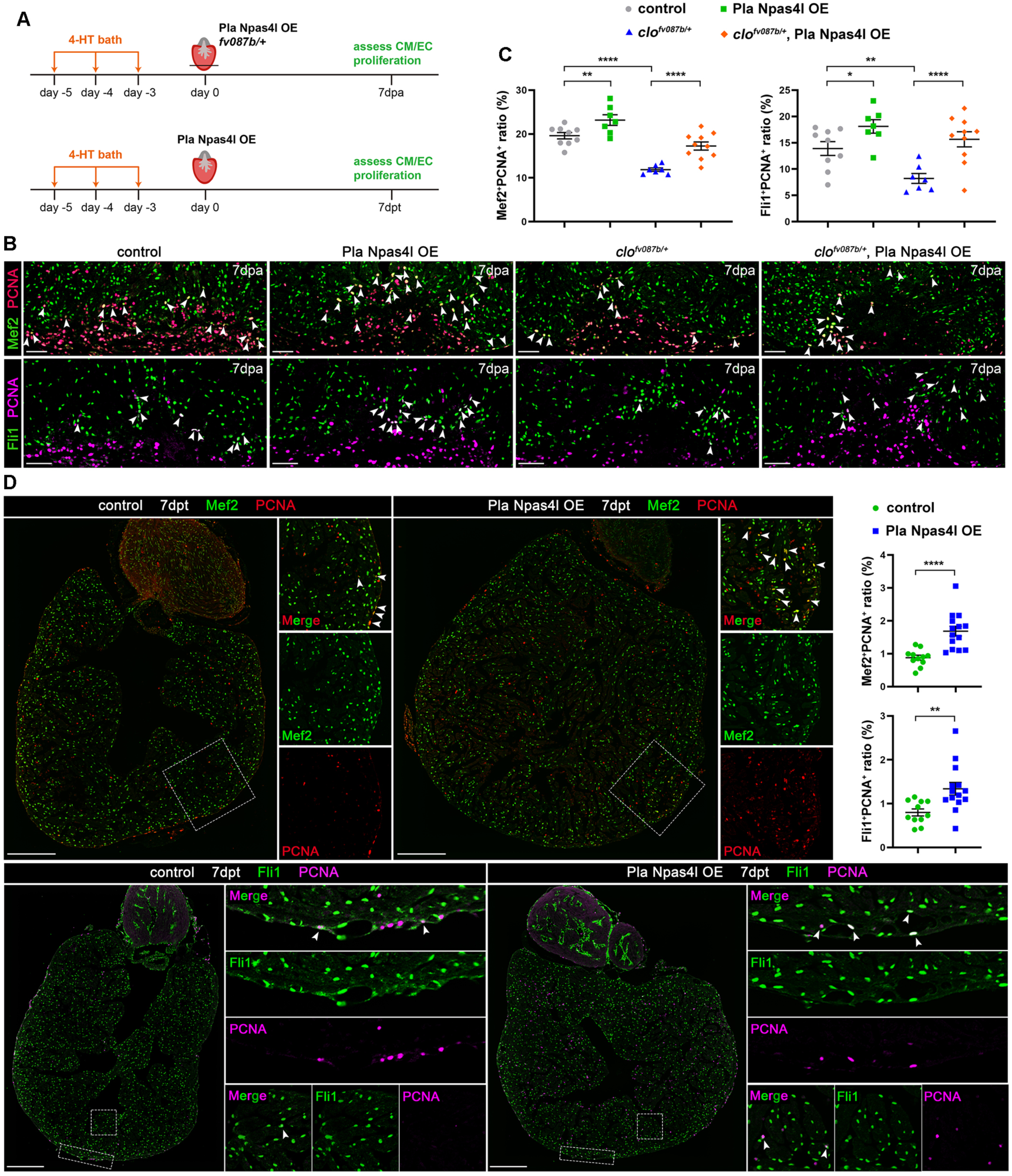
Platelet *npas4l* is required and sufficient for CM and EC proliferation. **A** Schematic diagram of experimental design to explore the function of platelet *npas4l* over-expression. **B, C** Representative immunofluorescence images (**B**) and quantification (**C**) of PCNA-positive CMs and ECs in *CD41:Cre-ERT2* (control, n=9), *CD41:Cre-ERT2; ubi:loxp-stop-loxp-npas4l-HA* (Pla Npas4l OE, n=7), *CD41:Cre-ERT2; clo^fv087b/+^* (*clo^fv087b/+^*, n=7) and *CD41:Cre-ERT2; ubi:loxp-stop-loxp-npas4l-HA; clo^fv087b/+^* (*clo^fv087b/+^*, Pla Npas4l OE, n=10) hearts at 7 dpa after 4-HT treatment. Arrowheads indicate proliferating CMs or ECs. Data are the mean ± SEM.; *p <0.05; **p<0.01; ****p <0.001; one-way ANOVA with LSD test. Scale bars, 50 μm. **D** Representative immunofluorescence images and quantification of PCNA-positive CMs and ECs in *CD41:Cre-ERT2* (control, n=11) and *CD41:Cre-ERT2; ubi:loxp-eGFP-stop-loxp-npas4l* (Pla Npas4l OE, n=14) hearts (uninjured) at 7 days post 4-HT treatment. Insets are high magnification of boxed areas. Arrowheads indicate proliferating CMs or ECs. Data are the mean ± SEM.; **p <0.01; ****p<0.001; unpaired, two-tailed Student’s *t* test. Scale bars, 200 μm.

## Discussion

### Novel Npas4l function in heart regeneration

Npas4l is well-known as being the earliest transcription factor for the specification of endothelial and hematopoietic lineages during zebrafish embryogenesis (Reischauer et al., 2016; Stainier et al., 1995). Previous research has established its critical function in determining endothelial and hematopoietic cell identity, and have identified its target genes during vascular development (Capon et al., 2022; Marass et al., 2019; Mattonet et al., 2022). However, its function in adult organ regeneration has never been addressed. In this work, we present, for the first time, Npas4l function in zebrafish heart regeneration as a platelet transcription factor that is required and sufficient for CM proliferation and regeneration. Haploinsufficiency of *npas4l* disrupted heart regeneration and impaired the injury-induced proliferation of CMs and ECs. Interestingly, being quite different from its expression pattern in zebrafish embryos, we found that *npas4l* was enriched in *gata1a^+^* blood cells, particularly *itga2b^+^*platelets in adult hearts, without being evident expression in endothelial cells. This striking expression difference between embryos and adult hearts suggested that Npas4l may function through different mechanisms during heart regeneration. It is well understood that cardiomyogenic factors like Neuregulin1 (Nrg1), YAP, miRNA-199a-3p, and Klf1, as well as the small molecular cocktail 5SM are sufficient to promote quiescent CMs into the cell-cycle and division in zebrafish and mouse heart regeneration (Bersell et al., 2009; Du et al., 2022; Eulalio et al., 2012; Gemberling et al., 2015; Monroe et al., 2019; Ogawa et al., 2021). Similarly, overexpression of *npas4l* in platelets, driven by either platelet-specific *CD41* or erythrocyte/platelet-specific *gata1a* promoter, promoted CM and EC proliferation in either uninjured or injured adult zebrafish hearts, thus revealing the first platelet signaling for inducing the proliferation of CMs and ECs in zebrafish or any animal systems. While this study focuses on the function of Npas4l in platelets, its potential role in erythrocytes cannot be entirely ruled out. Our scRNA-seq data showed that *npas4l* expression overlaps well with *gata1a* expression; this was in most of the platelets and a small part of erythrocytes revealed by quantitative RT-qPCR results. And we noted no evident ligand expression and interaction of erythrocytes with CMs and ECs, further supporting the conclusion that Npas4l function in platelets rather than erythrocytes. Most importantly, platelet-specific overexpression of *npas4l* driven by *CD41* promoter, like the *gata1a* promoter, was able to rescue regenerative defects in *cloche* mutant hearts as well as was sufficient to promote quiescent CMs into the cell-cycle. Together, this work presents the first evidence that *npas4l,* as a platelet factor, is required and sufficient for zebrafish heart regeneration.

### scRNA-seq data cover major cell types present in regenerating zebrafish hearts

By constructing a single-cell transcriptomic atlas of zebrafish heart regeneration, we acquired >60,000 cells covering the major cardiac cell types. Around half of these cells were CMs, presenting much more comprehensive coverage of CMs than previous reports and thus providing a rich source to address CM heterogeneity during heart regeneration (Honkoop et al., 2019; Ma et al., 2021). And we clustered CMs into 11 subsets and obtained some clues about CM3 and CM9, which were potential proliferative CM subgroups regulated by platelet Npas4l signaling. Similarly, we identified EC2 and EC7 as proliferative EC subgroups modulated by platelet Npas4l. By the global analysis of the ligand expression in cell populations, we noted that platelets, smooth muscle cells, and macrophages were the most secretory populations in cell-cell communications. Importantly, these cell-cell signaling transductions in *cloche* heterozygote mutants were prominently reduced during heart regeneration, and Npas4l regulated the expression of a cluster of ligands in platelets to achieve platelet interactions with other cell types during heart regeneration. Thus, this comprehensive scRNA-seq analysis of adult hearts results in the discovery of platelet *npas4l* function in heart regeneration and provides the field a rich resource for addressing cell-cell interactions and related signaling pathways during heart regeneration.

### Novel platelet function in heart regeneration

Platelets are derived from megakaryocytes, which are small and lack nuclei in mammals. For a long time, platelets were thought to modulate the inflammatory environment for tissue repair (Mezger et al., 2019). A recent study suggested that platelet activation regulates the infarct size in the heart and brain by ischemia-reperfusion injury (Rohlfing et al., 2022). In this work, we discovered a pro-regenerative function of platelets during zebrafish heart regeneration. Decreasing platelets by either NTR-based deletion or *mpl* mutants severely impaired zebrafish heart regeneration. Interestingly, a dynamic response of platelets to heart injury occurred even in a non-bleeding heart injury model by NTR-based cardiomyocyte ablation, thus suggesting that platelet activation not only serves for bleeding-arrest but also has a pro-regenerative function. For platelet activation mechanisms, our cell-cell interaction analysis revealed no thrombopoietin (THPO)/THPO receptor (MPL) expression from scRNA-seq data, while we noted an evident erythropoietin (EPO)/EPO receptor (EPOR) expression between platelets and CMs or ECs. This clue indicates that after heart injury, the EPO-EPOR pathway is preferred for activating thrombocytopoiesis (Lin et al., 2022). In this case, injured CMs and/or ECs release EPO that interacts with EPOR-expressing platelets, thus achieving platelet recruitment to the injury site for heart regeneration. This may explain the dynamic platelet response to heart injury. On the other hand, we noted significant expression of the ligands HBEGF, BMP6, and VEGFC in platelets that interacted with CMs and/or ECs. Except for VEGFC, which is well known for coronary vessel regeneration (El-Sammak et al., 2022), much remains unknown about how HBEGF and BMP6 function during heart regeneration. HBEGF is broadly expressed in various organs, and is reported to have a protective function in liver and diabetic wounds (Johnson and Wang, 2015; Takemura et al., 2013). During tissue repair and regeneration, HBEGF promotes cellular proliferation, migration, adhesion, and differentiation (Dao et al., 2018). And therapeutic delivery of HBEGF accelerates cutaneous wound healing (Johnson and Wang, 2013). Together with the data in this work, the above results suggest that HBEGF may also have pro-regenerative effects on zebrafish heart regeneration. BMP6 is a key regulator in iron homeostasis and bone formation in development. And endocardial BMP6 has been reported to influence cardiac EC proliferation during zebrafish heart regeneration (Fang et al., 2020). In this study, we report that platelet BMP6 acts downstream of Npas4l to mediate platelet-CM interactions during heart regeneration. Over-expression of BMP6 rescued impaired CM proliferation in *clo^fv087b/+^*hearts after injury, indicating that platelet BMP6 is required for CM proliferation. Other signals mediating Npas4l-induced CM and EC proliferation need to be further investigated. Together, our data support the hypothesis that platelets house various pro-regenerative secretory factors essential for heart regeneration, and platelet BMP6 is the downstream effector of Npas4l for fine-tuning CM proliferation and regeneration.

### Limitations of the Study

In this study, to explore the potential of *npas4l* in inducing mature cardiomyocytes into the cell-cycle, we ectopically overexpressed *npas4l* in erythrocytes/platelets in uninjured hearts, and found the increased numbers of PCNA^+^ cardiomyocytes after Npas4l overexpression, which were visualized by co-staining with cardiomyocyte marker Mef2. In addition, overexpressed *npas4l* in erythrocytes/platelets also promoted newly-formed coronary vessels and endothelial cell proliferation. However, the proliferation index showed a 1%-2% increase in Npas4l OE hearts compared with control hearts by using the proliferative CM or EC numbers to divide the total CM or EC numbers on the whole ventricle. These data thus suggested the effect of Npas4l on CM and EC proliferation might be small in uninjured hearts; on the other hand, due to the relatively low efficiency of *npas4l* overexpression, we only detected a 0.5-fold induction of *npas4l* mRNA no matter in the whole ventricles or isolated erythrocytes/platelets, which might account for small effect of Npas4l on CM or EC proliferation. Therefore, our data suggests a role of *npas4l* in inducing quiescent CM/EC into cell-cycle, but future studies are needed to thoroughly decipher the competence and mechanisms of *npas4l* on promoting cardiomyocyte proliferation in uninjured hearts. Furthermore, this work is restricted to study *npas4l* function in zebrafish heart regeneration, and it will be important to address whether it also play a conservative role in mammalian heart regeneration in the future.

## Acknowledgements

The authors thank Prof. Mark C. Fishman (Harvard University) and Prof. IC Bruce (Guest Professor at the College of Future Technology, Peking University) for commenting on and revising the manuscript; and members of Dr. Jing-Wei Xiong’s and Dr. Jianbin Wang’s laboratories for helpful discussions and technical assistance. The authors thank Dr. Yiyue Zhang at South-China University of Science and Technology (Guangzhou, China) for kindly providing *Tg(CD41:eGFP)^la2^* and *mpl^smu3^*zebrafish lines and Dr. Wenqing Zhang at South-China University of Science and Technology (Guangzhou, China) for kindly providing the *CD41:dsRed* construct. The authors thank the National Center for Protein Sciences at Peking University, particularly Dr. Liying Du at the Flow Cytometry Core for technical help on the Beckman Coulter MoFlo XDP and Dr. Guilan Li for technical help on single-cell RNA-seq library construction; and Drs. Liqin Fu, Siying Qin and Chunyan Shan at the Optical Imaging Platform of the Core Facility, School of Life Sciences and National Biomedical Imaging Center, Peking University for assistance with the Zeiss LSM710, LSM980 and Nikon LSM AXR. This work is supported by grants from the National Key R&D Program of China (2023YFA1800600 and 2018YFA0800501); the National Natural Science Foundation of China (32230032, 31730061, 81870198 and T2225005); and the Ministry of Science and Technology of the People’s Republic of China (2018YFA0800200). JH is supported by a postdoctoral fellowship from Nanchang University and an internal fund from the School of Basic Medical Sciences, Nanchang University.

## Author contributions

JH and CX designed and performed most of the experiments, and analyzed data; JH wrote the manuscript and CX contributed to revising the manuscript. YS designed the experiments on scRNA-seq, bulk RNA-seq and CUT&TAG, performed bioinformatics analysis of the data, and contributed to writing the manuscript. JS and YH contributed to performing scRNA-seq and bulk RNA-seq experiments and data analysis. YS and XM contributed to generating *bmp6* mutant and several transgenic zebrafish lines. QZ, SQ, and YX contributed to developing a protocol for dissociating zebrafish cardiac cells. YGC, XZ and JW supervised this work, designed experiments, analyzed data, and contributed to revising the manuscript. JWX conceived and supervised this work, designed experiments, analyzed data, and wrote the manuscript.

## RESOURCE AVAILABILITY

### Lead contact

Further information and requests for resources and reagents should be directed to and will be fulfilled by the Lead Contact, Jing-Wei Xiong, Ph.D.

### Materials availability

All unique/stable reagents generated in this study are available from the lead contact with a completed Materials Transfer Agreement.

### Data and code availability

Sequencing data that support the findings of this study were deposited into the GEO with the accession code GSE224125. Only public software and custom code were used to generate or process datasets in this work. The software version, references, and parameters are listed in the Key Resources Table and Method Details. Any additional information required to reanalyze the data reported in this paper is available from the lead contact upon request.

## EXPERIMENTAL MODEL AND SUBJECT DETAILS

### Zebrafish lines and maintenance

All zebrafish in this study were raised and handled according to a zebrafish protocol (IMM-XiongJW-3) approved by the Institutional Animal Care and Use Committee at Peking University, which is fully accredited by the Association for Assessment and Accreditation of Laboratory Animal Care International. Wild-type, mutant, and transgenic zebrafish of the outbred Tuebingen and AB strains of both males and females (4 to 10 months old) were used in this study. The transgenic and mutant zebrafish lines used were as follows: *clo^m39/+^* (Reischauer et al., 2016), *clo^fv087b/+^* (Xiong et al., 2008), *bmp6^pku422^*, *Tg(CD41:ECFP-NTR)^pku414^*, *Tg(gata1a:Cre-ERT2)^pku391^*,*Tg(CD41:Cre-ERT2)^pku417^*, *Tg(ubi:loxp-eGFP-stop-loxp-npas4l)^pku394^*, *Tg(ubi:loxp-stop-loxp-npas4l-HA)^pku419^*, *Tg(ubi:loxp-stop-loxp-bmp6)^pku420^*, *Tg(myl7:ECFP-NTR)^pku360^*(Han et al., 2014), *Tg(coro1a:eGFP)^hkz04t^* (Li and Hu, 2012), *Tg(gata1a:dsRed)^sd2^* (Traver et al., 2003), *Tg(CD41:eGFP)^la2^*(Lin et al., 2005), *Tg(kdrl:eGFP)^s843^* (Jin et al., 2005), *Tg(myl7:cypher-eGFP)^pku365^* (Xiao et al., 2016), *Tg(tcf21:dsRed2)^pd37^*(Kikuchi et al., 2011), *Tg(myl7:nucDsRed)^zf3348^* (Xie et al., 2020), and *Tg(gata4:eGFP)^ae1^* (Heicklen-Klein and Evans, 2004). The mutant zebrafish *mpl^smu3^* (Lin et al., 2017) were kindly provided by Dr. Yiyue Zhang. Zebrafish were housed at a density of <4 fish per liter. Water temperature was maintained at 28 °C. Ventricular resection was performed as described previously (Poss et al., 2002).

### Cell lines

HEK293T cell line (ATCC, SCSP-502) was cultured in DMEM (Cytiva, SH30022.01) with 10% fetal bone serum (Yeason, 40130ES76) and Penicillin-Streptomycin-Gentamicin Solution (Solarbio, P1410) at 37° C in a 5% CO_2_ humidified incubator.

## METHOD DETAILS

### Drug treatment

For metronidazole-mediated ablation of CMs, adult zebrafish were bathed in 1 mM metronidazole (Sigma, M1547) for 7 days before washout. For metronidazole-mediated ablation of platelets, adult zebrafish were bathed in 1 mM metronidazole the day before heart amputation. After heart injury, the zebrafish were bathed in 1mM metronidazole daily before harvesting the hearts. For 4-hydroxytamoxifen-mediated *npas4l* overexpression, adult zebrafish were bathed in 5 μM 4-hydroxytamoxifen (Sigma, H7904) for 10-12 h before washout. This cycle was repeated daily for total three times. For *bmp6* overexpression, the adult zebrafish received a heat shock in 39 °C water for 30 min. After cooling down to room temperature for 1 h, the zebrafish were bathed in 5 μM 4-hydroxytamoxifen for 10-12 h before washout. This cycle was repeated daily for total three times. All the zebrafish were maintained at a density of 4 zebrafish per 150 ml water for drug bathing. For BrdU tracing assays, BrdU (Sigma, B5002) was dissolved in DMSO to make a 100 mM stock solution. Adult zebrafish were anesthetized in E3 medium containing 0.4% tricaine and injected intrapericardially with 20 μl BrdU (8 mM in PBS) daily for three days before harvesting hearts. For BMP inhibitor treatment, LDN-193189 (MCE, HY-12071) and K02288 (MCE, HY-12278) were dissolved in DMSO to make 10 mM stock solutions. Adult zebrafish were anesthetized in E3 medium containing 0.4% tricaine and injected intrapericardially with 20 μl LDN-193189 (50 μM in PBS), K02288 (50 μM in PBS) or vehicle (0.5% DMSO in PBS) daily for three days before harvesting hearts.

### Generation of Tg(CD41:ECFP-NTR)^pku414^, Tg(CD41:Cre-ERT2)^pku417^, Tg(gata1a:Cre-ERT2)^pku391^, Tg(ubi:loxp-eGFP-stop-loxp-npas4l)^pku394^, Tg(ubi:loxp-stop-loxp-npas4l-HA)^pku419^, and Tg(ubi:loxp-stop-loxp-bmp6)^pku420^ transgenic zebrafish lines

*CD41:ECFP-NTR* was generated by replacing the *dsRed* sequence from *CD41:dsRed* construct which contains a 6-kb *CD41* promoter sequence, with the *ECFP-NTR* sequence. The *CD41:dsRed* construct was a gift from Dr. Wenqing Zhang, and the *ECFP-NTR* sequence was kindly provided by Dr. Ling-Fei Luo (Curado et al., 2007). *CD41:Cre-ERT2* was generated by replacing the *ECFP-NTR* sequence from the *CD41:ECFP-NTR* construct with the *Cre-ERT2* sequence. *gata1a:Cre-ERT2* was generated by replacing the *GFP* sequence from Gata1-GM2 vector which contains an 8.1-kb *gata1a* promoter sequence, with the *Cre-ERT2* sequence. The Gata1-GM2 vector was a gift from Dr. Shuo Lin (Long et al., 1997). *The Cre-ERT2* sequence was PCR-amplified from *myl7:Cre-ERT2* plasmid (Xiao et al., 2016). *ubi:loxp-eGFP-stop-loxp-npas4l* was generated by modifying the *ubi:loxp-DsRed-stop-loxp-eGFP* construct. which was kindly provided by Dr. C. Geoffrey Burns (Mosimann et al., 2011). The *DsRed* and e*GFP* sequences were replaced by *eGFP* and *npas4l* coding sequences, respectively. The *eGFP* sequence was PCR-amplified from *ubi:loxp-DsRed-stop-loxp-eGFP* plasmid and the *npas4l* coding sequence was synthesized by Beijing Ruiboxingke Biotechnology Co. according to the published sequence (Reischauer et al., 2016). *ubi:loxp-stop-loxp-npas4l-HA* was generated by modifying *ubi:loxp-eGFP-stop-loxp-npas4l* plasmid. The *eGFP* and *npas4l* sequences were replaced by 9 repeats of polyA signal and 6 repeats of HA tagged *npas4l* sequences, respectively. *ubi:loxp-stop-loxp-bmp6* was generated by replacing *npas4l-HA* sequence with *bmp6* coding sequence. The *bmp6* coding sequence was PCR-amplified from cDNA extracted from 24-hpf embryos. Each of the above entire cassettes was cloned into the Tol2 backbone and co-injected with transposase mRNA into one-cell stage embryos for making transgenic zebrafish (Kawakami et al., 2004).

### Generation of *bmp6^pku422^* mutant zebrafish line

The *bmp6^pku422^* zebrafish line was generated by using CRISPR/Cas9 technology with the following guide RNA sequences (5’-3’, PAM sequences in bold): GGGAGCTGAGCAGGGTTGGC**TGG** (exon 3 of *bmp6*) and GGCGTTGTTTGCCTGTAGAC**CGG** (exon 4 of *bmp6*). The protocol was described previously (Chang et al., 2013). In brief, *Cas9* mRNA (300 ng/μl) and gRNA (20ng/μl each) were co-injected into one-cell stage, wild-type embryos. The embryos were raised up and PCR-genotyping was used to identify the deletion mutants using the following primers (5’-3’): bmp6-genotype-F: GCAATACAATGCACTTGGGTCATGG; bmp6-genotype-R: CCACTCTTTAGTTGCTGTGCATACAC.

### Generation of anti-Npas4l polyclonal antibody

The peptide antigen of zebrafish Npas4l 395-647 amino acids tagged by GST was cloned, expressed, and purified by Huabio Co. (Hangzhou, China). Immunization was performed on mice and rabbits by the Laboratory Animal Center, Peking University and the serum was collected as polyclonal antibody.

### Cell transfection

To overexpress zebrafish Npas4l protein *in vitro*, HEK293T cells were transfected with pcDNA3.1-npas4l-flag-myc vector with Fugen HD transfection reagents (Promega, 2311) according to manufacturer’s instructions. pcDNA3.1-npas4l-flag-myc vector was generated by inserting the flag- and myc-tagged zebrafish *npas4l* coding sequence into pcDNA3.1 vector backbone.

### Adult zebrafish heart dissociation

For each sample, 15 adult ventricles were collected and placed in ice-cold perfusion buffer (in mM: 130 NaCl, 5 KCl, 0.5 NaH_2_PO_4_, 10 HEPES, 10 glucose, 10 2,3-butanedionemonoxime, 10 taurine, 5 MgCl_2_, pH adjusted to 7.8 with NaOH). The ventricles were gently chopped, transferred, and digested in digestion buffer (10 mg/ml protease, 30 μg/ml DNase I in perfusion buffer) for 3 h at 4 °C with constant gentle agitation using a rotator. The digestion was then neutralized with 10% FBS (fetal bovine serum) and the disassociated cells were filtered through a 100 μm-strainer to remove large clumps and pelleted at 500 × *g* for 10 min at 4 °C. The pellet was re-suspended in Hank’s balance salt solution (HBSS).

### Single-cell RNA-seqencing (scRNA-seq)

The dissociated cells were stained with Hoechst 33342 (1:300 diluted in HBSS), passed through a 70 μm-strainer, and loaded on a Beckman Coulter MoFlo XDP system to remove cell debris and sort for single cells. The sorted single cells were pelleted at 1000 × *g* for 5 min at 4 °C. The pellet was re-suspended in HBSS. Cell viability and concentration were measured on a Countstar Rigel system (ALIT Life Science). Samples with >85% cell viability were qualified for single-cell loading. Around 12,000 cells per sample were loaded into a 10x Genomics Chromium chip. Single-cell RNA-seq libraries were generated using Chromium Next GEM Single Cell 3’ Kit v3.1 (10x Genomics, 1000269) according to the manufacturer’s instructions. Quality control of the cDNA and library was performed on an Agilent 4150 TapeStation. The concentrations of cDNA and library were measured by a Qubit dsDNA high sensitivity assay (Invitrogen, Q33230). The sequencing was performed on an Illumina Novaseq PE150 platform operated by Peking Novogene Co.

### Single-cell RNA-seq data processing

The Cell Ranger (v3.0.2) pipeline (10x Genomics) was used to demultiplex the cellular barcodes and align reads to the GRCz11 zebrafish reference genome. The output filtered gene expression matrices were loaded and analyzed using the Seurat package (v3.1.4) (Butler et al., 2018). Potential doublet cells were detected and filtered using the R package DoubletFinder (v2.0.2) (McGinnis et al., 2019). Low-quality cells were removed from the datasets based on the following criteria: (1) the number of detected genes ≤200 or ≥4,000, the percentage of mitochondrial genes≥25 for non-CMs; and (2) the number of detected genes ≤200 or ≥4,000 for CMs. The mitochondrial content was not filtered due to the nature of CMs containing a high mitochondrial density (Gladka et al., 2018; Suryawanshi et al., 2020). Then, gene expression matrices were normalized by the NormalizeData function and highly variable genes were calculated using the FindVariableFeatures function.

### Cell clustering and annotation

To integrate cells into a shared space from different samples, we applied the integration methods described at https://satijalab.org/seurat/v3.0/integration.html (Stuart et al., 2019). In brief, we selected 2,000 features for integration using the SelectIntegrationFeatures function, identified ‘anchors’ between individual datasets with the FindIntegrationAnchors function, and integrated cells with identified ‘anchors’ using the IntegrateData function. Next, gene expression matrices were scaled with a linear transformation by the ScaleData function. Then, we applied principal component analysis (PCA) by the RunPCA function to achieve linear dimensional reduction after data scaling. After that, the ElbowPlot and JackStrawPlot functions were used to identify the dimensionality of the integrated datasets. Finally, we clustered cells using the FindNeighbors and FindClusters functions and performed nonlinear dimensional reduction with the RunTSNE function. To annotate cell types, the FindAllMarkers function was used to identify cluster-specific marker genes for each of the identified clusters, and the clusters were then classified and annotated based on the expression of canonical markers. Next, we applied a second round of clustering to further characterize subpopulations of CMs, ECs, and platelets. To avoid unexpected noise, mitochondrial genes and ribosome biogenesis genes were removed (Gladka et al., 2018; Luo et al., 2022; Suryawanshi et al., 2020; Xue et al., 2022). The integration, scaling, PCA, and clustering were applied as described above.

### Gene set variation analysis (GSVA)

The pathways used here are 50 hallmark pathways, GO terms, and KEGG pathways described in the molecular signature database (Subramanian et al., 2005). To obtain these pathways for zebrafish, we used the R package msigdbr (v7.0.1). Each cluster analyzed was downsampled to 200 cells because of the low cell number of some clusters. To assign pathway activity estimates to individual cells, we applied GSVA using standard settings, as implemented in the GSVA R package (v1.35.7) (Hanzelmann et al., 2013). The differential activities of pathways between clusters were calculated using the Limma R package (v3.38.3) (Ritchie et al., 2015). Significantly disturbed pathways were identified with a Benjamini-Hochberg-corrected P value of ≤0.05.

### Cell–cell communication analysis

To investigate potential interactions across different cell types, cell–cell communication was analyzed using CellPhoneDB (v2.1.4) (Efremova et al., 2020) with default parameters. In brief, a log2-normalized count matrix was subsampled into 300 cells per cluster. Then, we used the biomaRt R package (v2.38.0) (Durinck et al., 2009) to convert the genes of zebrafish to its orthologs in humans, due to the lack of interaction databases for zebrafish.

### RNA extraction and quantitative RT-PCR

For RNA extraction from adult zebrafish hearts, 10 ventricles were pooled to extract RNA for each sample. The total RNA was extracted using TRIzol reagent (Invitrogen, 15596026). RNA purity and concentration were determined using NanoDrop2000. 500 ng of total RNA was reverse-transcribed to cDNA using PrimeScript RT Reagent Kit (Takara, RR037A). RNA extraction and cDNA generation of sorted *CD41^+^* or *gata1a^+^*cells were performed using the SMART-seq2 method (Picelli et al., 2014) with modifications. In brief, ventricular dissection and cardiac cell dissociation were carried out as described above on *Tg(CD41:eGFP)* and *Tg(gata1a:dsRed)* hearts. Either 6,000 eGFP^+^ or dsRed^+^ cells were sorted into lysis buffer (0.5% Triton X-100, 1 U/μl RNase inhibitor, 2.5 μM Oligo dT primer, 2.5 mM dNTP) on a Beckman Coulter MoFlo XDP system. After vortex and incubation for 3 min at 72℃, total mRNA was reverse-transcribed to cDNA using SuperScript II Reverse Transcriptase (Invitrogen, 18064014). cDNA was then amplified using KAPA HiFi HotStart ReadyMix (Huaruikang, kk2602) and purified using VAHTS DNA Clean Beads (Vazyme, N411). cDNA concentration was measured using the Qubit dsDNA high sensitivity assay. Real-time quantitative PCR was carried out in triplicate using ChamQ SYBR Color qPCR Master Mix (Vazyme, Q411) on a Roche LightCycler96 system. To compensate for variations in RNA input and efficiency of reverse transcription, *rpl13a* was used for normalization. The primers used in this work were: *gapdh*-F: GATACACGGAGCACCAGGTT and *gapdh*-R: GCCATCAGGTCACATACACG; *rpl13a*-F: TCTGGAGGACTGTAAGAGGTATGC and *rpl13a*-R: AGACGCACAATCTTGAGAGCAG; *npas4l*-F: GCCACAGCAGACGCAGAGAT and *npas4l*-R: ACACTGAGGAGAGGCAGAGGAG; *bmp6*-F: GAACCGCAATCGCTCCAGTAGTC and *bmp6*-R: AGCCTTCAGGTGCAATAATCCAGTC; *vegfc*-F: GTCAACCTCTCAACCGCACCAA and *vegfc*-R: GCCATTGTCCGTTAGCATTCAGC; *admb*-F: CTAACCAACCAACCAAGACGCTCT and *admb*-R: CTGTCATGTCGCTCTCGCTTCTC; *apln*-F: ACACGCACACCACTACAGTATATCA and *apln*-R: GCTTCCCACCTCCTCAATCTCTTT; *tfpia*-F: GGAACGCCAACAACTTCAAGACCAT and *tfpia*-R: ATCTCCTCTCAGAGCCGTCACAATC; and *thbs1b*-F: CCTCAATCCACACCATCCTGACTC and *thbs1b*-R: TTGAACCACCACACCAAGACCTT; *bmp2a*-F: GTGAACGCAGAGCAGGTTAGCA and *bmp2a*-R: GGATGGAGGTCAGGTTGAAGAGGA; *bmp2b*-F: ACGCTTGCTCAATATGTTCGGATT and *bmp2b*-R: CTCGCCAGGAATGGAGGTAAGG; *bmp7a*-F: TGCTGGACTCACGAGTGGTATGG and *bmp7a*-R: GCGGAGGTGGATTCCTGATGCT; *bmp7b*-F: GAGACCTTGGATGGCAGGATTGGA and *bmp7b*-R: CGGAGATGGCGTGGAGTTGTGT.

### Bulk RNA-sequencing

RNA was extracted and cDNA was generated from sorted *CD41^+^*or *gata1a^+^* cells as above. 1 ng cDNA per sample was used to generate a library using the TruePrep DNA Library Prep Kit V2 for Illumina (Vazyme, TD503) according to the manufacturer’s instructions. In brief, 1 ng cDNA was fragmented using TruePrep Tagment Enzyme and then ligated with specific indexes. After amplification using TruePrep Amplify Enzyme, the cDNA library was purified using VAHTS DNA Clean Beads. The cDNA concentration was measured using Qubit dsDNA high sensitivity assay. The sequencing was performed on an Illumina Novaseq PE150 platform operated by Peking Novogene Co.

### Bulk RNA-seq data processing

Raw sequencing data quality was analyzed using FastQC (v0.11.4) and processed by Trim Galore (v0.6.0) to remove adapters and low-quality reads. Clean reads were aligned to the GRCz11 zebrafish reference genome using HISAT2 (v2.1.0) (Kim et al., 2019). The read counts were generated with HTseq (0.6.1p1) (Temin, 1966). The genes needed to have 10 reads in at least one replicate for subsequent differential expression analysis. Differential gene expression was analyzed using the DESeq2 R package (v1.22.1) (Love et al., 2014). Genes with a p-value <0.05 and a log2 fold change >0.485 (fold change >1.4) were considered to represent significantly differentially expressed genes (DEGs). The pathway analysis of DEGs was performed using the Metascape webtool (www.metascape.org) (Zhou et al., 2019).

### ChIP-seq with CUT&TAG method

For each sample, 30 adult ventricles were collected and digested as described above with modifications. The ventricles were gently chopped, transferred, and digested in digestion buffer for 15 min at 4 °C with constant gentle agitation using a rotator. The digestion was then neutralized with 10% FBS (fetal bovine serum) and the dissociated cells were filtered through a 100 μm-strainer to remove large clumps and pelleted at 500 × *g* for 10 min at 4 °C. The pellet which mainly contained platelets and erythrocytes was re-suspended in Hank’s balance salt solution (HBSS). Then, around 2000,000 cells per sample were fixed in 0.2% formaldehyde for 2 min and neutralized by adding glycine to a final concentration 25 mM. The fixed cells were pelleted at 800 × *g* for 5 min at 4 °C, washed twice with PBS and loaded for CUT&TAG reaction using Hyperactive Universal CUT&TAG Assay Kit for Illumina Pro (Vazyme, TD904) and Mouse anti-HA Tag antibody (Santa Cruz, sc-7392) according to the manufacturer’s instructions. The concentrations of library were measured by a Qubit dsDNA high sensitivity assay. The sequencing was performed on a BGI DNBSEQ-T7 PE150 platform operated by Peking GenePlus Co. Raw sequencing data was processed as mentioned above. Clean reads were aligned to the GRCz11 zebrafish reference genome using the Bowtie 2 (v2.4.5) software (Langmead and Salzberg, 2012). The aligned BAM files were then sorted and deduplicated using the samtools (v1.15.1) and Picard (v2.1.0) to mitigate PCR artifacts (Danecek et al., 2021). The MACS2 (v2.2.7.1) was employed with parameters “-p 0.05” to call peaks (Zhang et al., 2008). The deepTools (v3.5.0) and ggcoverage (v1.4.0) were used to generate the heatmap and track plot separately (Ramirez et al., 2016; Song and Wang, 2023). The motif enrichment is performed using HOMER (v3.12) with default parameters (Heinz et al., 2010).

### Whole-mount RNAscope *in situ* hybridization in zebrafish embryos

Zebrafish embryos were collected and then fixed in 10% NBF (neutral buffered formalin, Solarbio, G2161) at room temperature for 24 h. The fixed embryos were washed with PBST (0.1% Tween-20 in PBS) and dehydrated. After drying for half an hour, the embryos were treated with Protease Plus (ACD, 322330) for 20 min at room temperature followed by washout with PBST. For *npas4l* hybridization assay, RNAscope 2.5 HD Reagent Kit-BROWN (ACD, 322300) was applied to visualize signals according to the manufacturer’s instructions with modifications. In brief, the RNAscope Probe-Dr-*npas4l* (ACD, 483481) probe was applied to the embryos for hybridization (overnight at 40℃). After washout with 0.2× SSCT, the embryos were post-fixed in 4% paraformaldehyde/PBS at room temperature for 10 min (Gross-Thebing et al., 2014). The embryos were incubated with Amp1-6 as described in the instructions and washed with 0.2× SSCT after each Amp treatment. The signals were visualized using DAB substrates provided in the kit mentioned above. After washout with water, dehydration by serial methanol dilution, clearing with benzyl benzoate (Sigma, B17700) and benzyl alcohol (Sigma. B16208) mixture (2 volumes of benzyl benzoate and 1 volume of benzyl alcohol), the embryos were fixed in neutral balsam (Solarbio, G8590) and photographed under a Leica DM5000B microscope.

### RNAscope *in situ* hybridization on fresh frozen heart sections

Zebrafish hearts were dissected, freshly embedded in Optimal Cutting Temperature (OCT) compound (Sakura, 4583), and stored at −80 °C. 7 μm-thick cryosections were cut on a Leica CM1950 Cryostat Microtome. RNAscope pre-treatment was performed using RNAscope H_2_O_2_ and Protease Reagent Kits (ACD, 322381) according to the manufacturer’s instructions. In brief, cryosections were fixed in pre-chilled 4% paraformaldehyde/PBS for 15 min at 4 °C. After dehydration and air drying for 5 min, the sections were treated with RNAscope hydrogen peroxide for 10 min at room temperature. After washout with water, the slides were incubated with RNAscope Protease IV for 8 min at room temperature and then washed with PBS. For *npas4l* sole-hybridization assays, RNAscope 2.5 HD Reagent Kit-BROWN (ACD, 322300) was applied to visualize signals according to the manufacturer’s instructions. For dual-hybridization assays, RNAscope 2.5 HD Duplex Reagent Kit (ACD, 322430) was used to visualize signals according to the manufacturer’s instructions. For *itga2b* sole-hybridization assays, RNAscope 2.5 HD Duplex Reagent Kit was applied to visualize signals according to the manufacturer’s instructions with modifications. In brief, the pre-treated slides were incubated with target probe and AMP1-6 as described in the instructions followed by signal visualization. For *bmp6* sole-hybridization assays, RNAscope 2.5 HD Duplex Reagent Kit was applied to visualize signals according to the manufacturer’s instructions with modifications. In brief, the pre-treated slides were incubated with target probe and AMP1-3, 8-10 as described in the instructions followed by signal visualization. All the above slides were co-stained with eosin (Solarbio, G1120) or Mayer’s hematoxylin (Solarbio, G1080) and mounted with Eukitt(R) quick-hardening mounting medium for microscopy (Sigma, A2303989). For *cre*-*itga2b* and *cre-gata1a* dual-hybridization co-stained with MHC immunofluorescence assays, the pre-treated slides were incubated with anti-MHC monoclonal antibody (eBioscience, 14-6503-82, 1:300 diluted in Co-Detection Antibody Diluent) overnight at 4 °C. The next day, the slides were washed with PBST and fixed in 10% neutral formalin fix solution (NBF) for 30min at room temperature. After washout with PBST, the RNAscope Multiplex Fluorescent Detection Reagents v2 was used to visualized the signals according to the manufacturer’s instructions. After signal visualization, the slides were incubated with secondary antibody (1:300 diluted in Co-Detection Antibody Diluent) for 1h at room temperature. After washout with PBST, the slides were mounted with fluorescence mounting medium with DAPI (ZSGB-BIO, ZLI-9557). For *npas4l* sole-hybridization with MHC immunofluorescence assays, RNAscope 2.5 HD Reagent Kit-BROWN was applied to visualize signals according to the manufacturer’s instructions with modifications. In brief, after visualization of RNAscope signals, the slides were blocked in blocking solution (1% Tween-20, 10% FBS, 1% DMSO in PBS) for 1 h at room temperature and incubated with anti-MHC monoclonal antibody (1:300 diluted in blocking solution) overnight at 4 °C. The next day, the slides were washed with PBS and incubated with secondary antibody (1:300 diluted in blocking solution) for 2 h at room temperature. After washout with PBS, the slides were mounted with fluorescence mounting medium with DAPI. The RNAscope probes used in this study were: RNAscope Probe-Dr-*npas4l* (ACD, 483481), RNAscope Probe-Cre (ACD, 312281), RNAscope Probe-Dr-*bmp6*-C1 (ACD, 1105911-C1), RNAscope Probe-Dr-*gata1a*-C2 (ACD, 473371-C2), RNAscope Probe-Dr-*itga2b*-C2 (ACD, 555601-C2), RNAscope Probe-Dr-*kdrl*-C2 (ACD, 416611-C2), RNAscope Probe-Dr-*postnb*-C2(ACD, 525381-C2), and RNAscope Probe-Dr-*tcf21*-C2 (ACD, 485341-C2). All images were captured under a Leica DM5000B and Nikon LSM AXR microscopes.

To quantify *npas4l* signals in injured hearts, 3 fields (0.2 × 0.15 mm) per heart within the wound area were acquired using a ×63 objective and the *npas4l^+^* areas were measured using ImageJ. To quantify *itga2b* signals in injured heart, *itga2b^+^* areas within the wound area were measured using ImageJ. The ratio of *itga2b^+^* area *versus* the length of the apical wound was calculated for each heart. To quantify *itga2b* signals in uninjured heart, *itga2b^+^* areas within the entire ventricle were measured using ImageJ. Sections with the largest wound area for apex resected hearts or with the largest ventricle size for uninjured hearts or hearts with genetic ablation of cardiomyocytes were analyzed.

### Immunofluorescence on adherent cultured cells

The cells were washed three times with pre-chilled PBS then fixed in 4% paraformaldehyde/PBS for 15 min at room temperature. After three washes with PBS, the cells were blocked with blocking solution (1% BSA, 0.1% Triton X-100 in PBS) for 1 h at room temperature then incubated with Rabbit anti-Npas4l antibody (1:300 diluted in blocking solution) and Mouse anti-Myc Tag (Engibody, AT0023, 1:300 diluted in blocking solution) at 4 °C overnight. The next day, the cells were washed three times with 1% BSA/PBST (0.02% Tween-20 in PBS) then incubated with donkey anti-rabbit Alexa Fluor 488 (Invitrogen, A21206, 1:300 diluted in blocking solution), goat anti-mouse Alexa Fluor 555 (Invitrogen, A21422, 1:300 diluted in blocking solution) for 2 h at room temperature. After three washes with PBST, the cells were mounted with fluorescence mounting medium with DAPI (ZSGB-BIO, ZLI-9557). All images were captured under Zeiss LSM710, LSM980 and Nikon LSM AXR confocal microscopes.

### Immunofluorescence on zebrafish heart sections

Zebrafish hearts were dissected and fixed in 4% paraformaldehyde/PBS for 1 h at room temperature. After dehydration, the hearts were embedded in paraffin using a Leica HistoCore Arcadia H-Heated Paraffin Embedding Station. 5 μm-thick paraffin sections were cut on a Leica RM2265 microtome. The paraffin sections were used for immunofluorescence as described previously (Han et al., 2014; Xiao et al., 2016). The primary antibodies used in this work were: Rabbit anti-Mef2c (Sigma, HPA005533, 1:200), Rabbit anti-Mef2a+Mef2c (Abcam, ab197070, 1:300), Rabbit anti-Fli1 (Abcam, ab133485, 1:300), Mouse anti-PCNA (Santa Cruz, sc-25280, 1:300), Mouse anti-PCNA (Santa Cruz, sc-56, 1:50), Rat anti-BrdU (Abcam, ab6326, 1:300), Rabbit anti-GFP (Invitrogen, A-11122, 1:300), Rabbit anti-Sarcomeric alpha Actinin (Abcam, ab68167, 1:300) and Mouse anti-myosin heavy-chain monoclonal antibody (eBioscience, 14-6503-82, 1:500). The secondary antibodies used were: donkey anti-rabbit Alexa Fluor 488 (Invitrogen, A21206, 1:300), goat anti-mouse Alexa Fluor 555 (Invitrogen, A21422, 1:300), donkey anti-mouse Alexa Fluor 488 (Invitrogen, A32766, 1:300) and goat anti-rat Alexa Fluor 555 (Invitrogen, A21434, 1:300). All images were captured under Zeiss LSM710, LSM980 and Nikon LSM AXR confocal microscopes.

To quantify CM or EC proliferation, Mef2^+^/PCNA^+^, Mef2^+^/BrdU^+^, Fli1^+^/PCNA^+^, or Fli1^+^/BrdU^+^ cells were manually counted and then were divided by the total numbers of Mef2^+^ or Fli1^+^ nuclei to get the proliferation index. For each ventricular resected heart, the cells within the area 100 μm from the wound edge were included in the analysis. For each uninjured heart or heart with genetic ablation of CMs, the cells within the entire ventricle were included in analysis. To quantify ECs, immune cells or platelets, Fli1^+^, *coro1a*:eGFP^+^, and *CD41*:eGFP^+^ areas within the wound were measured using ImageJ and then normalized by the length of the wound. Sections with the largest wound area for apex resected hearts or with the largest ventricle size for uninjured hearts or hearts with genetic ablation of cardiomyocytes were analyzed.

### Acid Fuchsin Orange-G staining (AFOG)

Paraffin sections were used for AFOG staining as described previously (Xiao et al., 2016). The slides were photographed under a Leica DM5000B microscope. To quantify regeneration level, the hearts were categorized into three levels of regeneration, not regenerated (severe), partially regenerated (partial), and fully regenerated (regenerated) according to the scar size. Serial sections for each heart were analyzed.

### Protein extraction and western blot

For protein extraction from adult zebrafish hearts, 10 ventricles were pooled to extract protein for each sample. The ventricles were homogenized in RIPA lysis buffer (Solarbio, R0010) with 1 mM PMSF (Sigma, 93482), PhosSTOP phosphatase inhibitor cocktail (Roche, 04906845001), and a protease inhibitor cocktail (Roche, 11873580001). The supernatant was collected for each protein sample. Then the protein was quantified using the BCA Protein Assay Kit (Solarbio, PC0020) and loaded with Prestained Protein Marker (Vazyme, MP102) for western blots as described previously (Zhou et al., 2022). The primary antibodies used were: Mouse anti-Npas4l (1:1000), Mouse anti-Myc Tag (Engibody, AT0023, 1:1000), Rabbit anti-Bmp6 (Abcam, ab155963, 1:3000), and Rabbit anti-β-actin (Huabio, ET1701-80, 1:3000). The secondary antibodies used were: goat anti-rabbit IgG-HRP (Easybio, BE0101, 1:3000) and goat anti-mouse IgG-HRP (Easybio, BE0102, 1:3000). Imaging was performed using the ChemiDoc MP Imaging System with Image Lab software (Bio-Rad). Quantification was performed using ImageJ and relative expression was normalized by β-actin.

### Statistical analysis

Data are presented as the mean ± standard error of the mean (SEM). The value of n is noted in the figures/figure legends and always stands for separate biological replicates. Statistical analysis was performed using IBM SPSS Statistics 20. Statistical differences between two samples were compared using unpaired, two-tailed Student’s *t* test. Comparative statistics among multiple samples were performed using one-way ANOVA with the LSD test. Categorical variables between two samples were analyzed using Pearson chi-square test. scRNA-seq data were analyzed using unpaired, two-tailed Fisher’s exact test, and P values were adjusted using the Benjamini-Hochberg method. A P value <0.05 was considered statistically significant and individual p values are noted in the figures/figure legends.

## Supplementary Figures and Figures Legends

**Supplementary Figures S1-S17**

**Supplementary Tables S1-S7**

