## Supplementary figures and legends for "Cloche/Npas4l is a pro-regenerative platelet factor during zebrafish heart regeneration"

### Supplementary Figures and Figure Legends

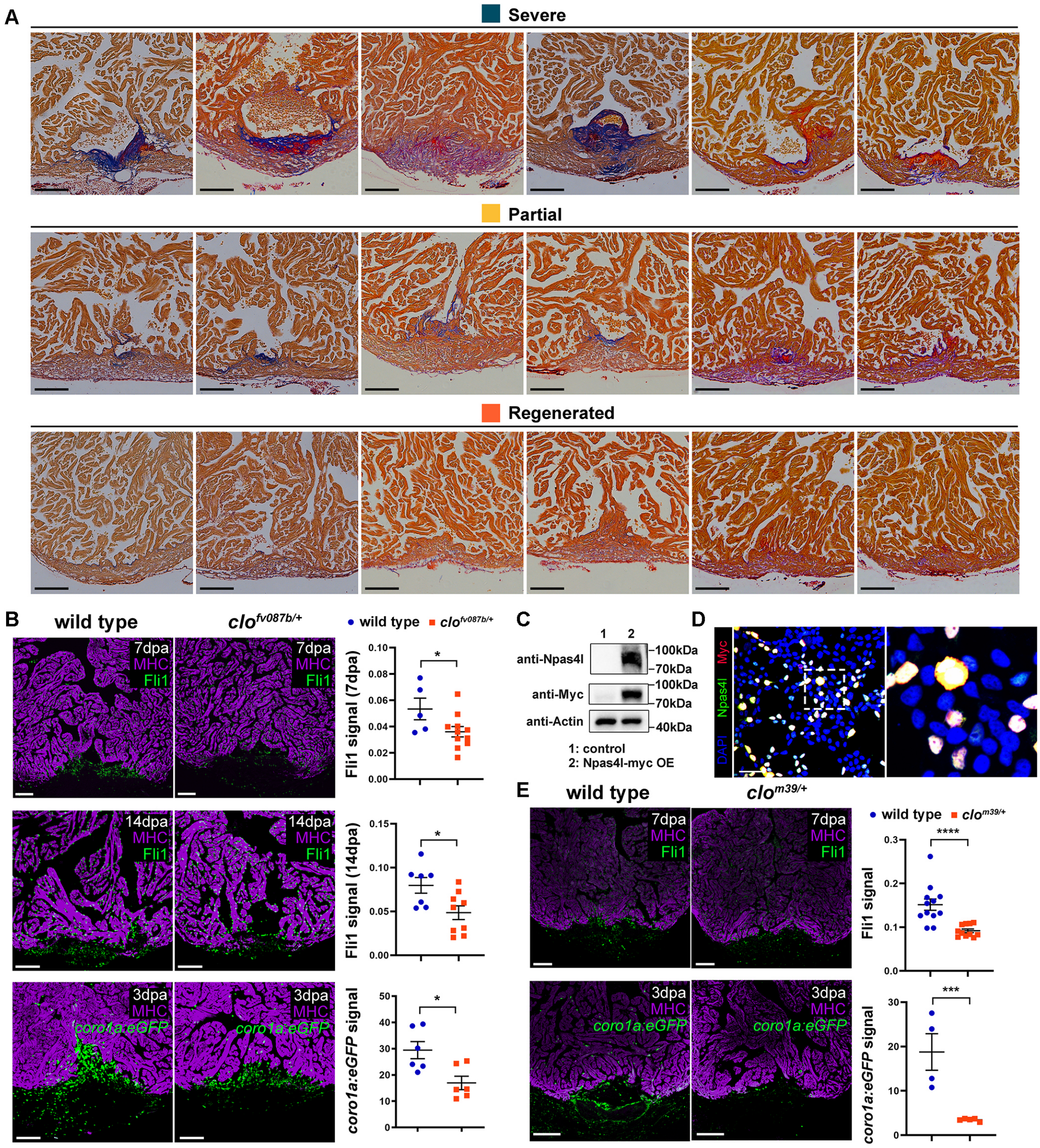


**Figure S1. Zebrafish anti-Npas4l antibody is successfully generated and haploinsufficiency of *npas4l* disrupts heart regeneration in both *cloche*$^{fv087b/+}$ and *cloche*$^{m39/+}$ mutants.**

**A** Representative images of AFOG staining, shown in Figure 1E-F, on not regenerated (severe), partially regenerated (partial), and fully regenerated (regenerated) hearts sections at 30 dpa. Scale bars, 100 μm. **B** Representative immunofluorescence images and quantification of Fli1-positive ECs or *coro1a:eGFP*-positive leukocytes co-stained with MHC antibody in wild-type sibling (n=5-7) and *clo*$^{fv087b/+}$ mutant (n=6-11) hearts at 3 dpa, 7 dpa and 14 dpa. Data are the mean ± SEM.; *p <0.05; unpaired, two-tailed Student's *t* test. Scale bars, 100 μm. **C** Western blot of Myc and Npas4l in control and myc-tagged Npas4l OE-293T cells, revealing the 92 kDa Npas4l band recognized by both anti-Npas4l and anti-Myc tag antibodies. Protein loading is normalized by β-actin. **D** Representative immunofluorescence images of Npas4l antibody co-stained with Myc tag antibody in Myc-tagged Npas4l OE-293T cells. The inset is high magnification of the boxed area. Scale bar, 50 μm. **E** Representative immunofluorescence images and quantification of Fli1-positive ECs or *coro1a:eGFP*-positive leukocytes co-stained with MHC antibody in wild-type sibling (n=4-12) and *clo*$^{m39/+}$ mutant (n=5-11) hearts at 3 dpa and 7 dpa. Data are the mean ± SEM.; ***p <0.005, ****p<0.001; unpaired, two-tailed Student's *t* test. Scale bars, 100 μm.

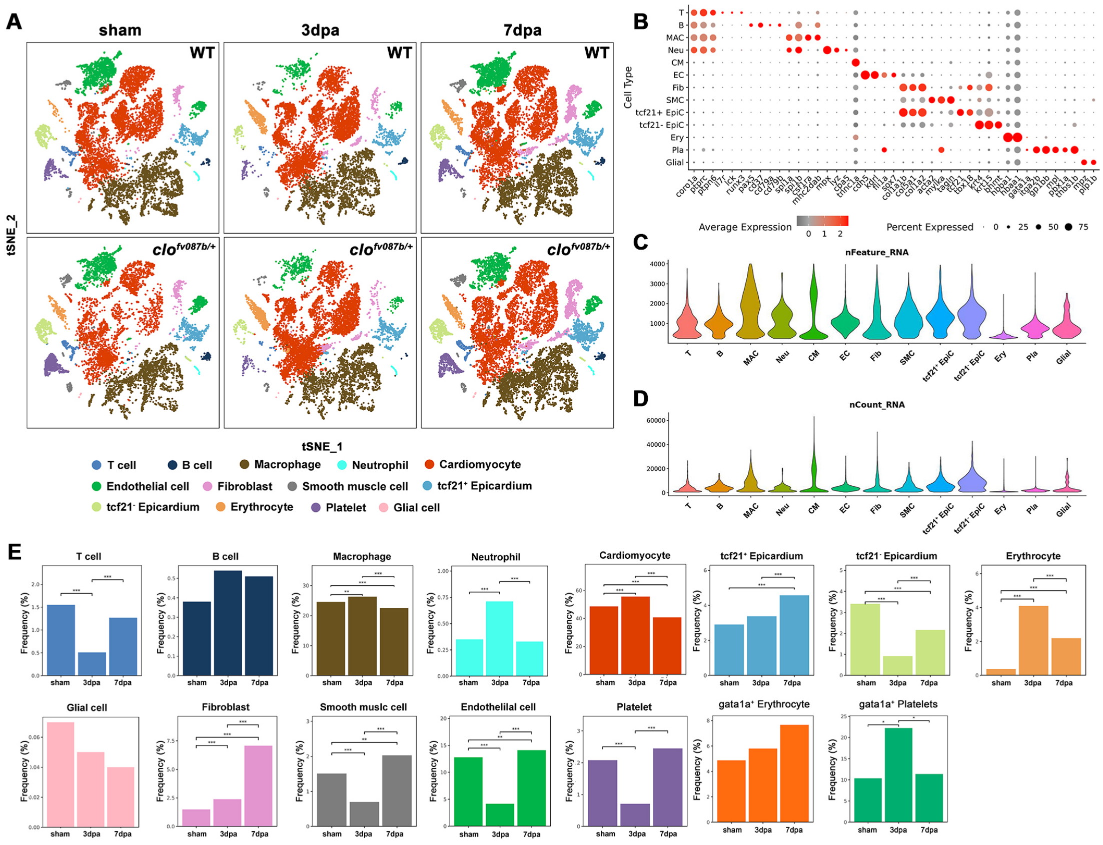


**Figure S2. Single-cell transcriptome atlas of wild-type sibling and *clo*$^{fv087b/+}$ mutant hearts.**

**A** t-SNE visualization of clusters from wild-type and *clo*$^{fv087b/+}$ hearts at sham, 3 dpa, and 7 dpa. **B** Dot plot showing marker gene expression in each cell cluster. **C, D** Violin plots depicting average nFeatures (gene numbers, **C**) and nCounts (UMI numbers, **D**) of each cell cluster. **E** Quantitative analysis of each cell cluster frequency from wild-type hearts at sham, 3 dpa, and 7 dpa. All differences Benjamini-Hochberg adjusted *p <0.05, **p <0.005, and ***p <0.001; two-tailed Fisher's exact test.

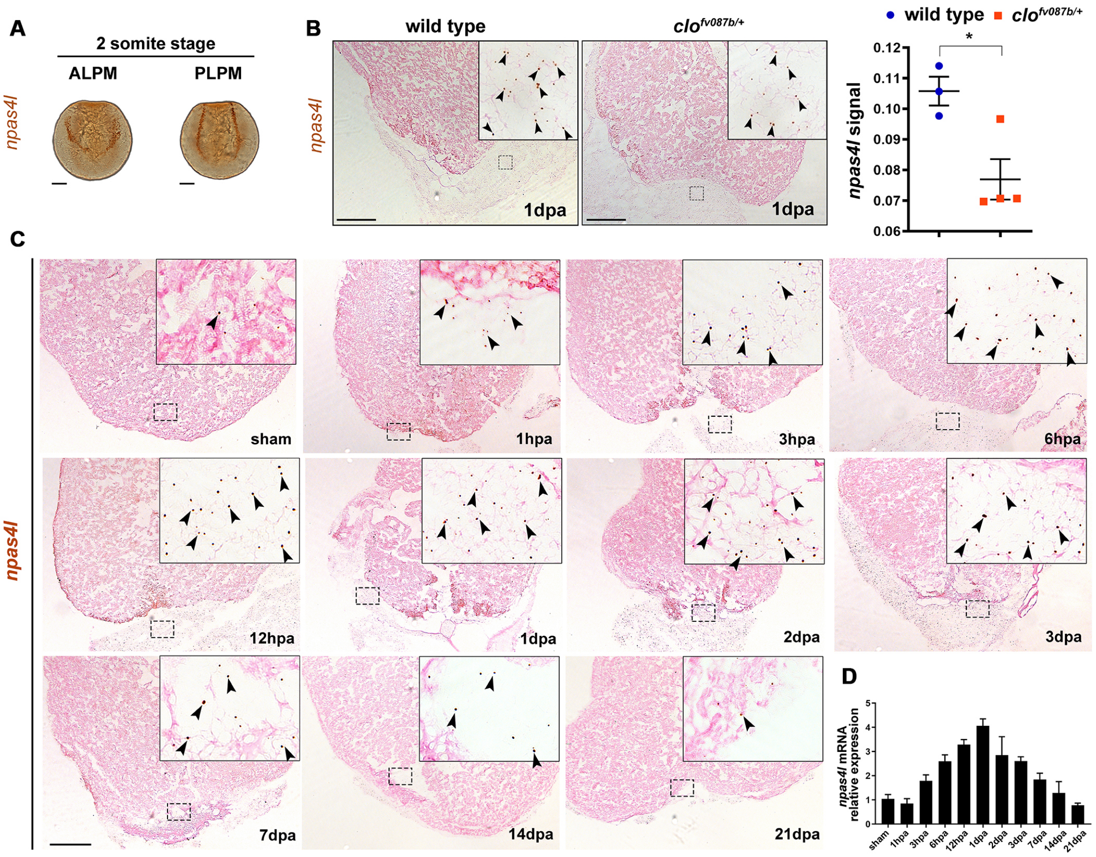


**Figure S3. Zebrafish *npas4l* is induced after ventricular resection.**

**A** Representative images of whole-mount RNAscope *in situ* hybridization with *npas4l* probes in 2-somite zebrafish embryos. Scale bars, 100 µm. **B** Representative images of RNAscope *in situ* hybridization with *npas4l* probes in wild-type and $clo^{fv087b/+}$ heart sections at 1 dpa. Insets are high magnification of boxed areas. Arrowheads indicate *npas4l* signals. n=3-4 hearts per group, 3 fields per heart. Data are the mean ± SEM.; ****p <0.001; unpaired, two-tailed Student's *t* test. Scale bars, 200 µm. **C** Representative images of RNAscope *in situ* hybridization with *npas4l* probe in zebrafish hearts after ventricular resection in the sham and from 1 h to 21 days post ventricular resection. Insets are high magnification of boxed areas. Arrowheads indicate *npas4l* signal. Scale bar, 200 µm. **D** Quantitative RT-PCR of *npas4l* mRNA expression normalized by *rpl13a* in zebrafish hearts after ventricular resection. n=3 technical replicates per group, 10 hearts pooled together per group. Data are the mean ± SEM.

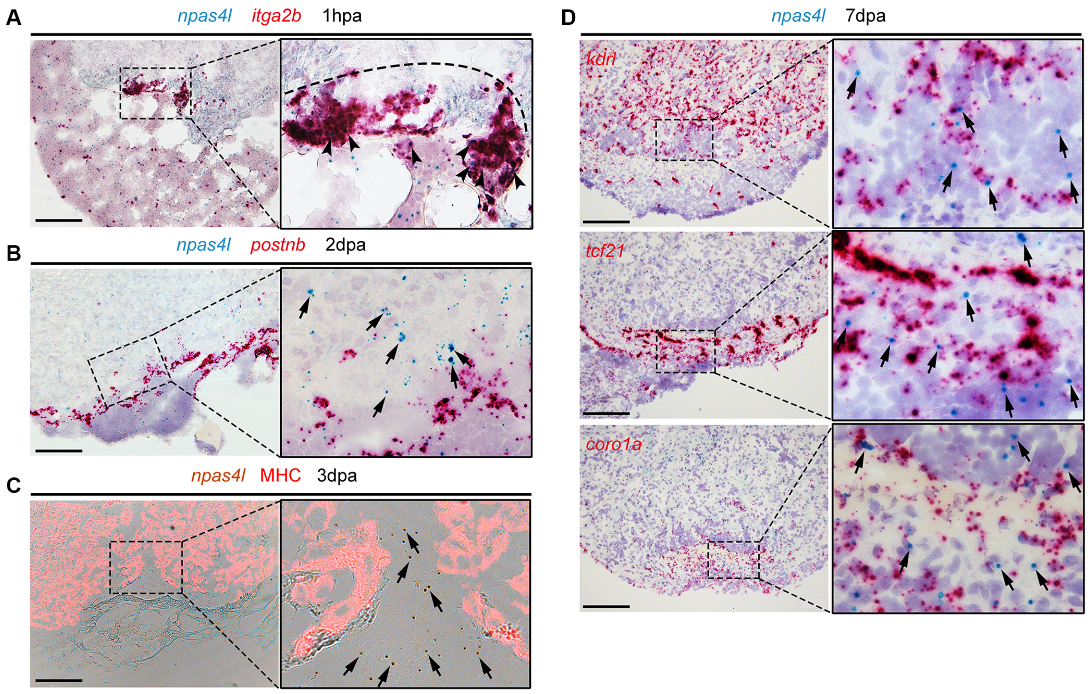


**Figure S4. Zebrafish *npas4l* is enriched in platelets at the wound edge after ventricular resection.**

**A** Representative images of RNAscope *in situ* hybridization with *npas4l* probe co-stained with *itga2b* probe in zebrafish hearts at 1 hpa. The inset is high magnification of the boxed area. Dashed lines delineate the wound edge. Arrowheads indicate *npas4l* and *itga2b* double positive signals. Scale bar, 100 μm. **B** Representative images of RNAscope *in situ* hybridization with *npas4l* probes co-stained with *postnb* probes in zebrafish hearts at 2 dpa. The inset is high magnification of the boxed area. Arrows indicate *npas4l* signals. Scale bar, 100 μm. **C** Representative images of *npas4l* RNAscope *in situ* hybridization co-stained with MHC immunofluorescence in zebrafish hearts at 3 dpa. The inset is high magnification of the boxed area. Arrows indicate *npas4l* signals. Scale bar, 100 μm. **D** Representative images of RNAscope *in situ* hybridization with *npas4l* probe co-stained with either *kdrl*, *tcf21,* or *coro1a* in zebrafish hearts at 7 dpa. Insets are high magnification of boxed areas. Arrows indicate *npas4l* signals. Scale bars, 100 µm.

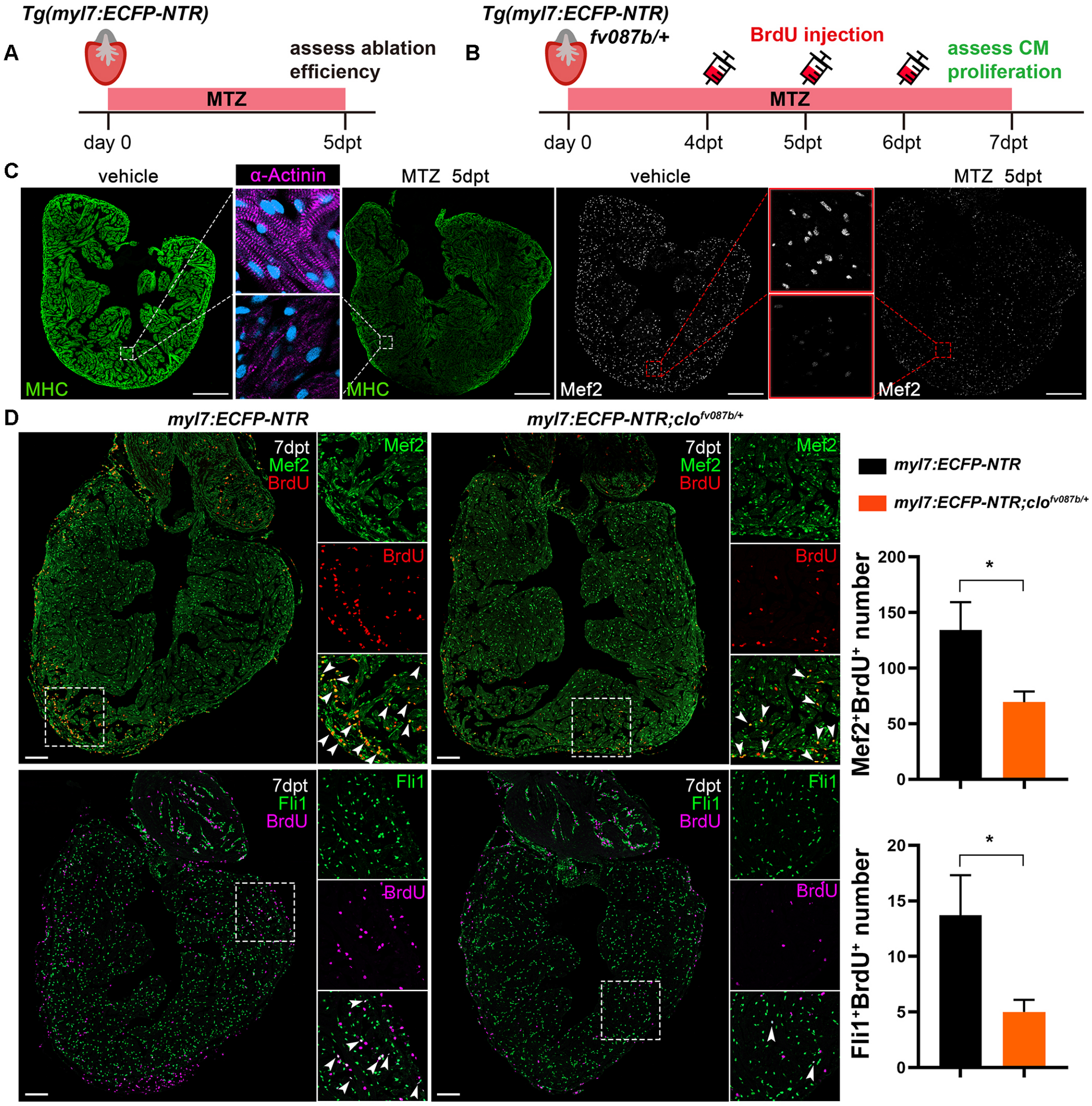


**Figure S5. Haploinsufficiency of *npas4l* impairs CM and EC proliferation in an NTR-mediated ablation injury model.**

**A** Schematic diagram of experimental design to assess NTR-mediated CM ablation efficiency. **B** Schematic diagram of experimental design to explore *npas4l* function in NTR-mediated ablation injury model. **C** Representative images of immunofluorescence of α-Actinin co-stained with MHC (left) and Mef2 (right) in vehicle (n=10) and MTZ (n=5) treated *Tg(myl7:ECFP-NTR)* hearts at 5 days post treatment. Insets are high magnification of boxed areas. Scale bars, 200 µm. **D**

Representative immunofluorescence images and quantification of BrdU-positive CMs and ECs in *myl7:ECFP-NTR* control and *myl7:ECFP-NTR:clo*$^{fv087b/+}$ mutant hearts at 7 days post MTZ treatment. Insets are high magnification of boxed areas. Anti-Mef2C and anti-Fli1 label CMs and ECs, respectively. Arrowheads indicate proliferating CMs or ECs. n=10 hearts per group. Data are the mean ± SEM.; *p <0.05; unpaired, two-tailed Student's *t* test. Scale bars, 100 μm.

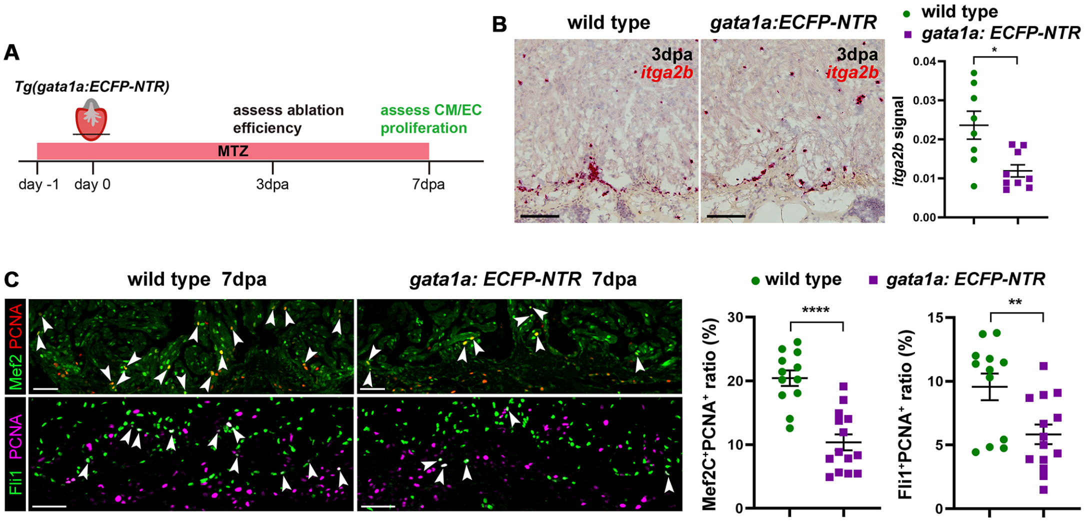


**Figure S6. Platelet ablation driven by the *gata1a* promoter disrupts zebrafish heart regeneration.**

**A** Schematic diagram of experimental design to explore platelet function during heart regeneration by platelet ablation. **B** Representative images of RNAscope *in situ* hybridization and quantification of *itga2b* in wild-type (n=8) and *gata1a:ECFP-NTR* hearts (n=9) at 3 dpa after MTZ treatment. Data are the mean ± SEM.; *p <0.05; unpaired, two-tailed Student's *t* test. Scale bars, 100 μm. **C** Representative immunofluorescence images and quantification of PCNA-positive CMs and ECs in wild-type sibling (n=12) and *gata1a:ECFP-NTR* (n=14) transgenic hearts at 7 dpa. Arrowheads indicate proliferating CMs or ECs. Data are the mean ± SEM.; **p <0.01; ****p <0.001; unpaired, two-tailed Student's *t* test. Scale bars, 50 μm.

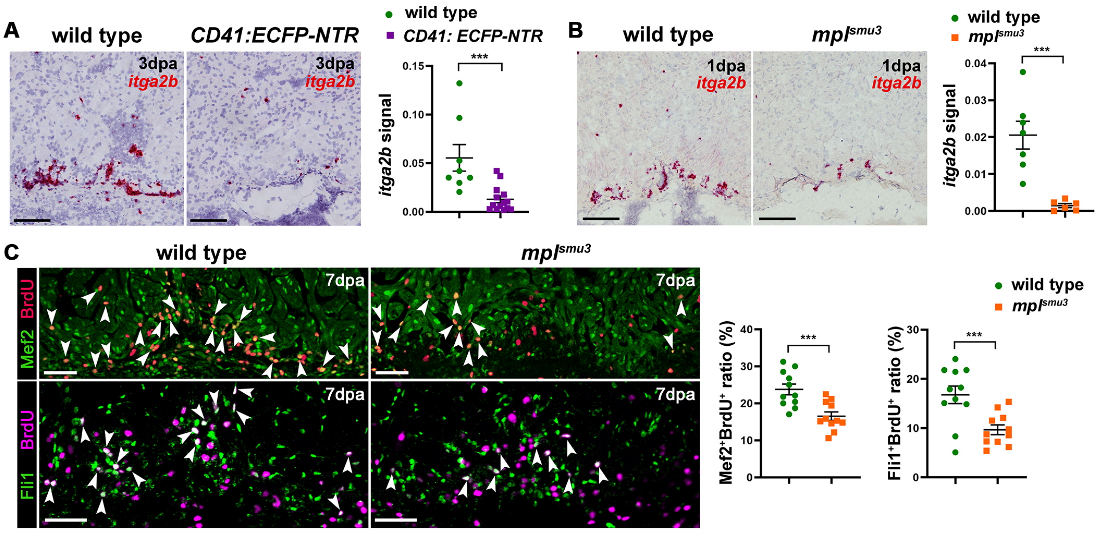


**Figure S7. Either platelet ablation driven by the CD41 promoter or platelet-deficient *mpl*$^{smu3}$ mutant disrupts zebrafish heart regeneration.**

**A** Representative images of RNAscope *in situ* hybridization and quantification of *itga2b* in wild-type (n=8) and *Tg(CD41:ECFP-NTR)* (n=14) hearts at 3 dpa after MTZ treatment. Data are the mean ± SEM.; ***p <0.005; unpaired, two-tailed Student's *t* test. Scale bars, 100 μm. **B** Representative images of RNAscope *in situ* hybridization and quantification of *itga2b* in wild-type control (n=7) and *mpl*$^{smu3}$ mutant hearts (n=6) at 1 dpa. Data are the mean ± SEM.; ***p <0.05; unpaired, two-tailed Student's *t* test. Scale bars, 100 μm. **C** Representative immunofluorescence images and quantification of BrdU-positive CMs and ECs in wild-type and *mpl*$^{smu3}$ mutant hearts at 7 dpa. Arrowheads indicate proliferating CMs or ECs. n=11 hearts per group. Data are the mean ± SEM.; ***p <0.005; unpaired, two-tailed Student's *t* test. Scale bars, 100 μm.

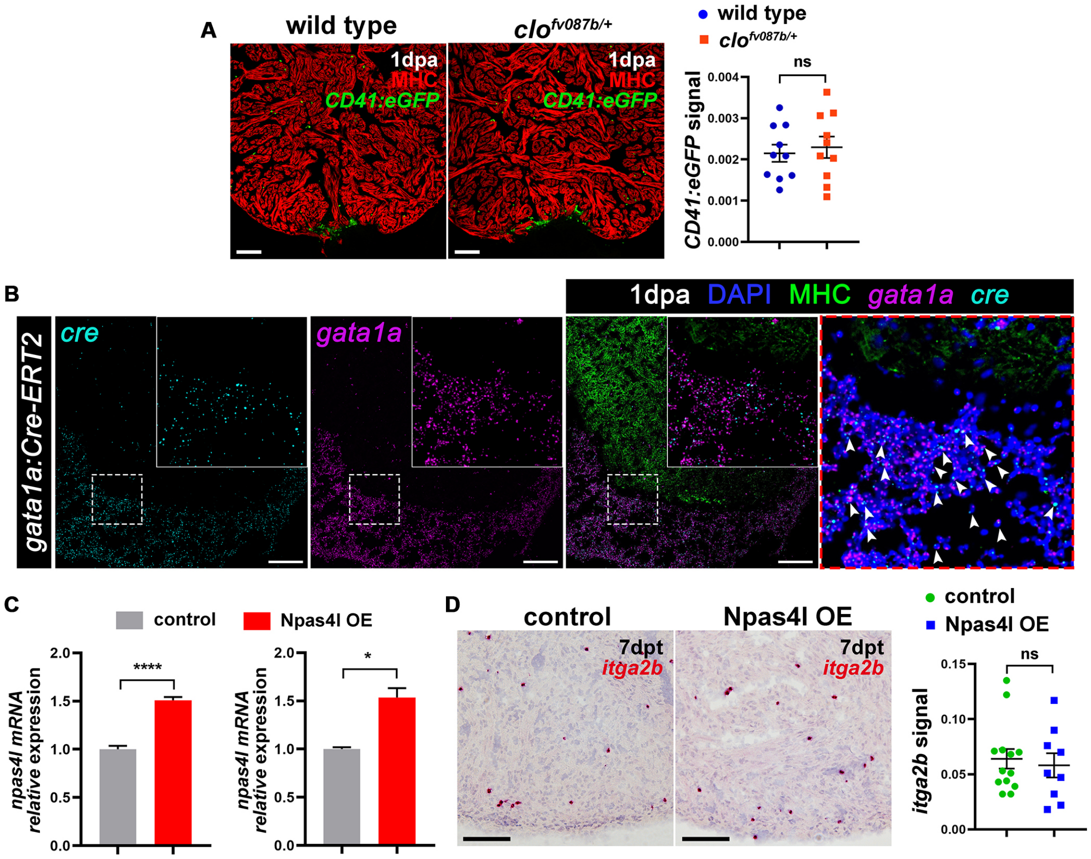


**Figure S8. Neither *npas4l* haploinsufficiency nor over-expression affects the numbers of platelets after injury.**

**A** Representative immunofluorescence images and quantification of *CD41:eGFP* co-stained with MHC antibody in wild-type and *clo*$^{fv087b/+}$ hearts at 1 dpa. n=10 hearts per group. Data are the mean ± SEM.; unpaired, two-tailed Student's *t* test. Scale bars, 100 µm. **B** Representative images of *cre-gata1a* RNAscope *in situ* hybridization co-stained MHC (labeling myocardium) on *Tg(gata1a:Cre-ERT2)* heart sections at 1dpa. The insets and red-outlined panel were the high magnifications of the boxed area. Arrowheads indicate the *cre* and *gata1a* double positive cells. Scale bars, 100 µm. **C** left: Quantitative RT-PCR of *npas4l* mRNA expression normalized by *rpl13a* in whole zebrafish ventricles at 3 days post 4-HT treatment. n=3 replicates per group, 10 hearts pooled together per group. Data are the mean ± SEM.; ****p <0.001; unpaired, two-tailed Student's *t* test. Right: Quantitative RT-PCR of *npas4l* mRNA expression normalized by *rpl13a* in isolated *gata1a*$^{+}$ platelets/erythrocytes at 7 days post 4-HT treatment. n=3 replicates per group and around 6000 cells pooled together per replicate. Data are the mean ± SEM.; *p <0.05; unpaired, two-tailed Student's *t* test. **D** Representative images of RNAscope *in situ* hybridization with *itga2b* probe in *Tg(ubi:loxp-eGFP-stop-loxp-npas4l)* (control, n=13) and *Tg(gata1a:Cre-ERT2; ubi:loxp-eGFP-stop-loxp-npas4l)* (Npas4l OE, n=9) hearts at 7 days post 4-HT treatment. Data are the mean ± SEM.; unpaired, two-tailed Student's *t* test. Scale bars, 100 μm.

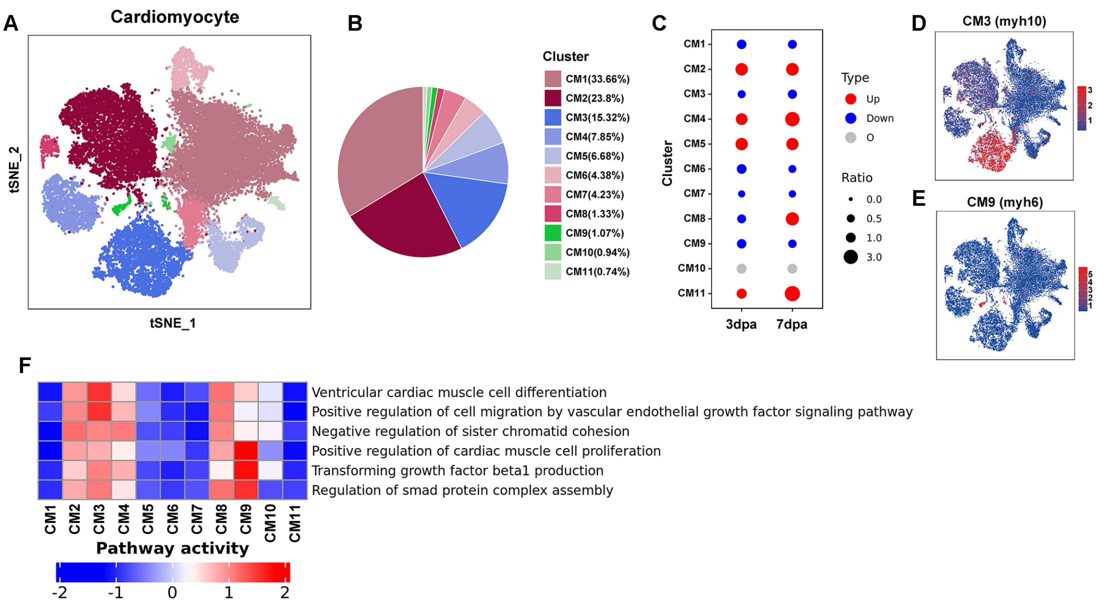


**Figure S9. scRNA-seq data reveals proliferating subclusters of cardiomyocytes.**

**A** t-SNE visualization of subclusters for cardiomyocytes from wild-type and *clo*$^{fv087b/+}$ hearts. **B** Pie plot showing proportion of each subcluster. **C** Dot plots showing that subclusters of cardiomyocytes increased (red) and decreased (blue) in *clo*$^{fv087b/+}$ hearts compared with wild-type sibling hearts at 3 dpa and 7 dpa (normalized by corresponding sham group). **D, E** Feature plots of CM3 (*myh10-positive*, **D**) and CM9 (*myh6-positive*, **E**) marker genes shown in panel A. **F** Heatmaps showing enriched gene set variation analysis (GSVA) of CM3 and CM9.

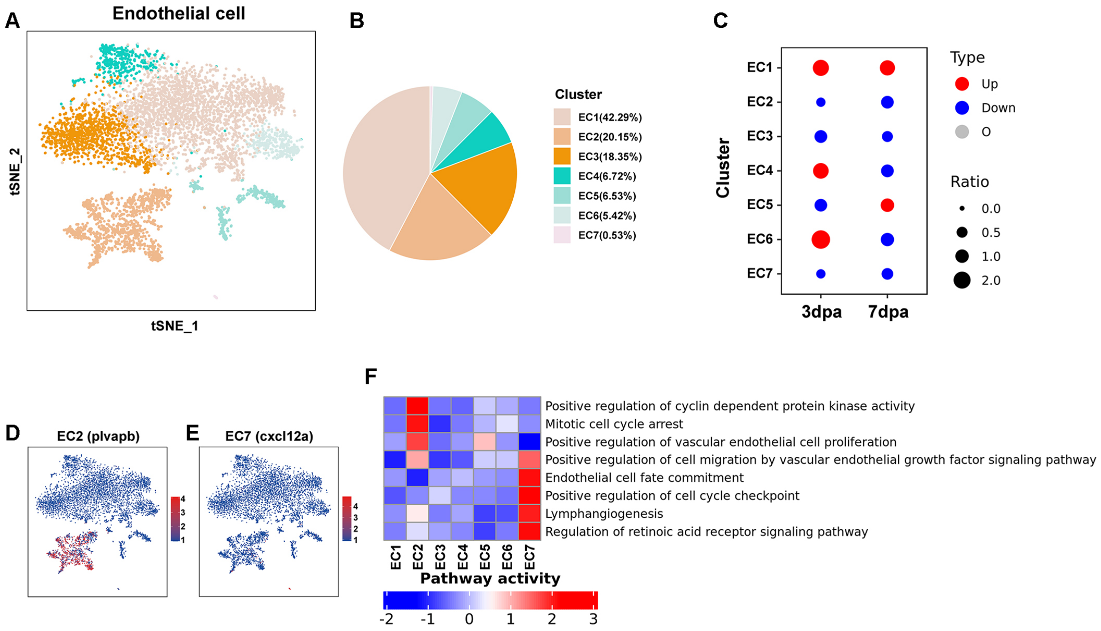


**Figure S10. scRNA-seq data reveals subclusters of endothelial cells.**

**A** t-SNE visualization of subclusters for endothelial cells from wild-type and *clo*$^{fv087b/+}$ hearts. **B** Pie plot showing proportion of each subcluster. **C** Dot plots showing that subclusters of endothelial cells increased (red) and decreased (blue) in *clo*$^{fv087b/+}$ hearts compared with wild-type sibling hearts at 3 dpa and 7 dpa (normalized by corresponding sham group). **D, E** Feature plots of EC2 (*plvapb-positive*, **D**) and EC7 (*cxcl12a-positive*, **E**) markers genes shown in panel A. **F** Heatmaps showing enriched GSVA of EC2 and EC7.

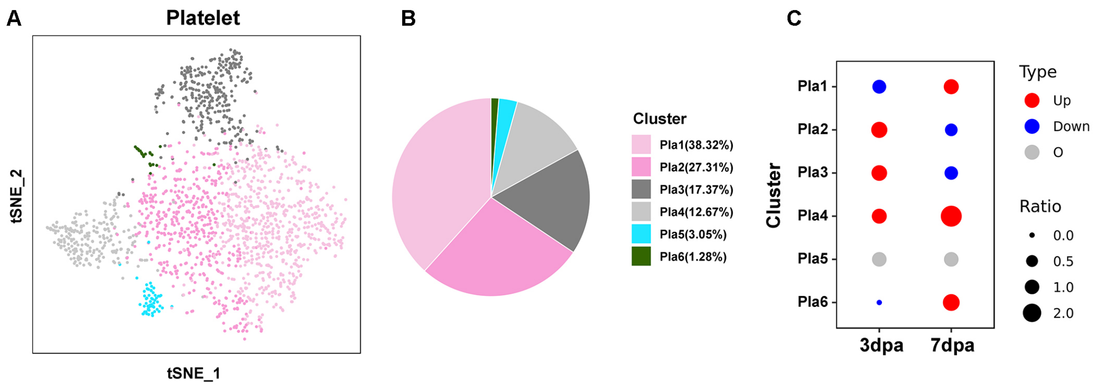


**Figure S11. scRNA-seq data reveals subclusters of platelets.**

**A** t-SNE visualization of subclusters for platelets from wild-type and *clo*$^{fv087b/+}$ hearts. **B** Pie plot showing proportion of each subcluster. **C** Dot plots showing that subclusters of platelets increased (red) and decreased (blue) in *clo*$^{fv087b/+}$ hearts compared with wild-type sibling hearts at 3 dpa and 7 dpa (normalized by corresponding sham group).

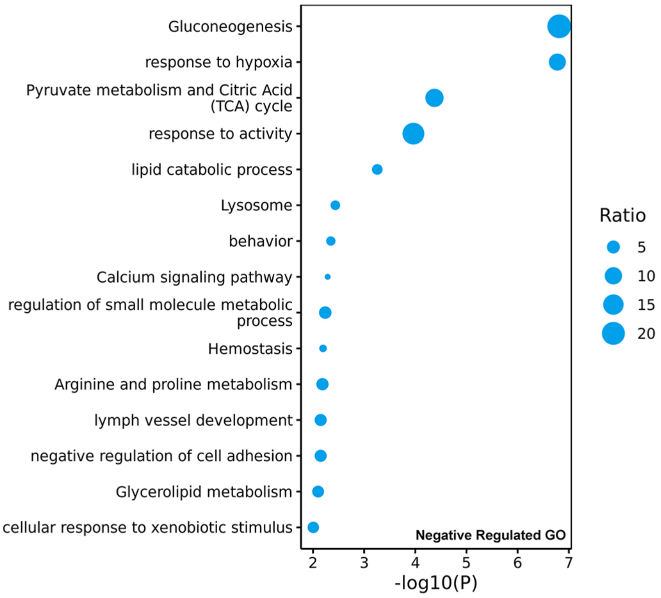


**Figure S12. Pathway analysis showing enriched terms of Npas4l negatively-regulated genes.**

Significantly enriched pathways of Npas4l negatively-regulated genes.

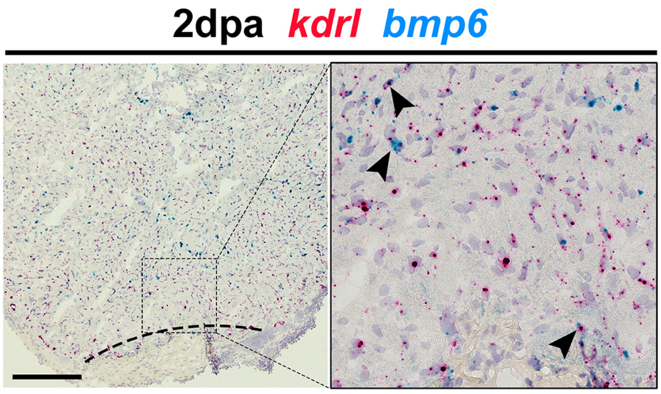


**Figure S13. *bmp6* was also expressed in endothelial cells within the border zone after injury.**

Representative images of RNAscope *in situ* hybridization using *bmp6* probe co-staining with *kdrl* probe in zebrafish hearts at 2 dpa. The inset is high magnification of the boxed area. Arrowheads indicate *bmp6*$^{+}$ and *kdrl*$^{+}$ overlapped signals. Dashed lines delineate the wound edge. Scale bar, 100 μm.

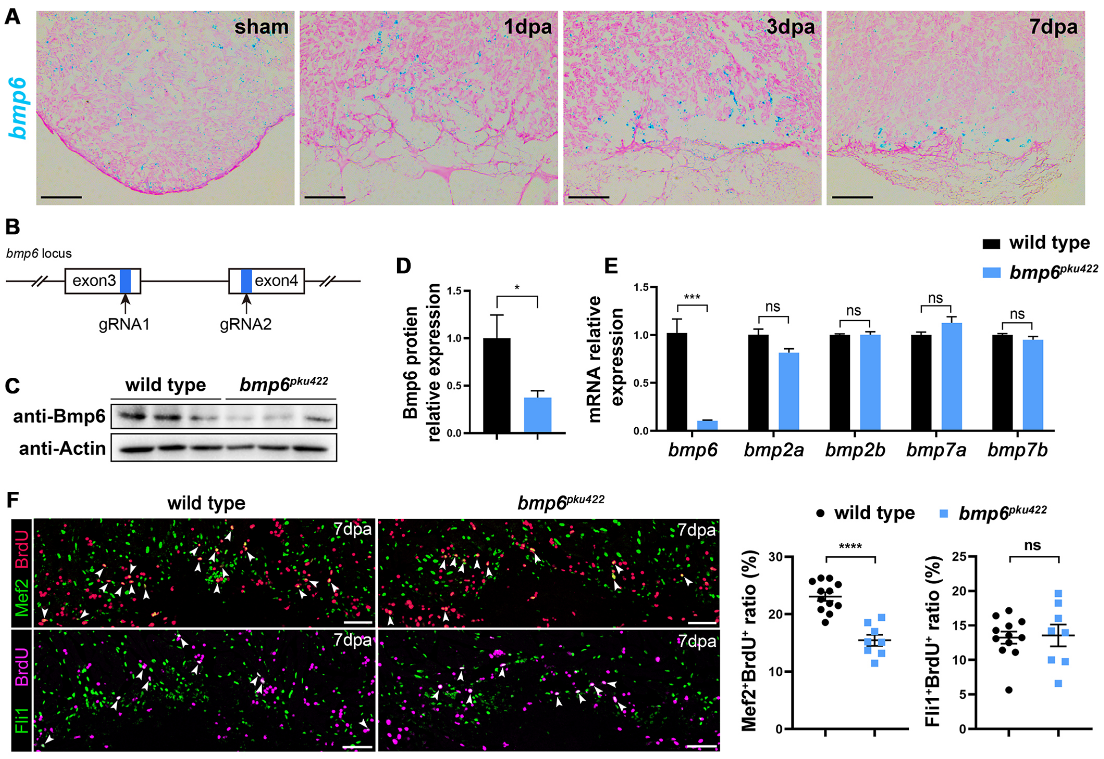


**Figure S14. *bmp6* was required for zebrafish heart regeneration.**

**A** Representative image of RNAscope *in situ* hybridization using *bmp6* probe in zebrafish hearts at sham, 1 dpa, 3 dpa and 7 dpa. n=3-6 hearts pre group. Scale bars, 100 µm. **B** Strategy to generate *bmp6* mutant zebrafish line. **C, D** Western Blot (**C**) and quantification (**D**) of *bmp6* expression in wild-type and *bmp6*$^{pku422}$ embryos at 24 hours post fertilization. Protein loading is normalized by β-actin. n=3 replicates, around 100 embryos pooled together per replicate. Data are the mean ± SEM.; *p <0.05; unpaired, two-tailed Student's *t* test. **E** Quantitative RT-PCR reveals expressions of *bmp6*, *bmp2a*, *bmp2b*, *bmp7a* and *bmp7b* in wild-type and *bmp6*$^{pku422}$ embryos at 24 hours post fertilization. mRNA loading is normalized by *rpl13a*. n=3 replicates, around 100 embryos pooled together per replicate. Data are the mean ± SEM.; ***p <0.005; unpaired, two-tailed Student's *t* test. **F** Representative immunofluorescence images and quantification of BrdU-positive CMs and ECs in wild-type (n=12) and *bmp6*$^{pku422}$ mutant (n=8) hearts at 7 dpa. Arrowheads indicate proliferating CMs or ECs. Data are the mean ± SEM.; ****p <0.001; unpaired, two-tailed Student's *t* test. Scale bars, 50 μm.

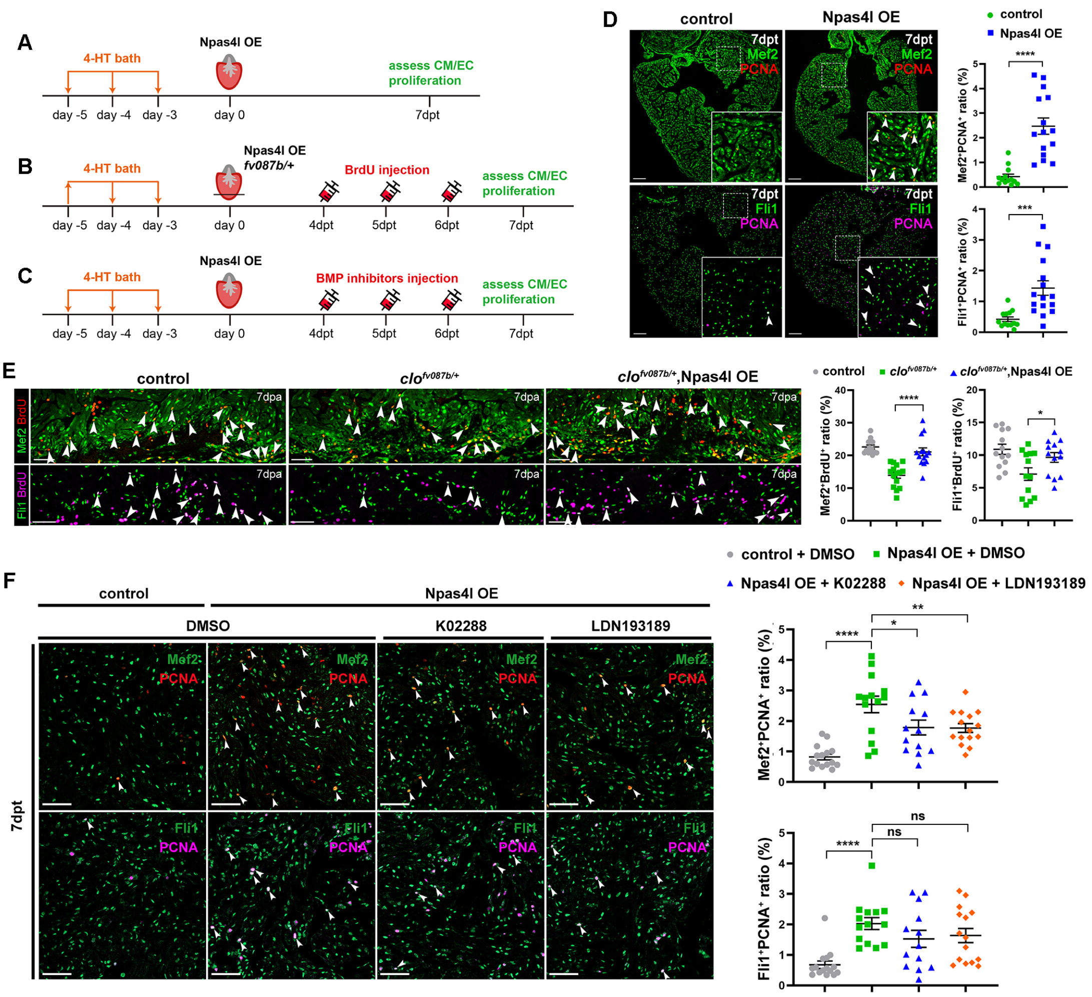


**Figure S15. Platelet *npas4l* is required and sufficient for CM and EC proliferation.**

**A** Schematic diagram of experimental design to explore the function of *npas4l* overexpression on uninjured hearts. **B** Schematic diagram of experimental design to explore the function of *npas4l* overexpression on rescuing *clo*$^{fv087b/+}$ phenotype. **C** Schematic diagram of experimental design to explore the function of BMP signal downstream to *npas4l*. **D** Representative immunofluorescence images and quantification of PCNA-positive CMs and ECs in *ubi:loxp-eGFP-stop-loxp-npas4l* (control: n=13) and *gata1a:Cre-ERT2; ubi:loxp-eGFP-stop-loxp-npas4l* (Npas4l OE: n=15-16) hearts at 7 days post 4-HT treatment (dpt). Insets are high magnification of boxed areas. Arrowheads indicate proliferating CMs or ECs. Data are the mean ± SEM.; ***p <0.005; ****p <0.001; unpaired, two-tailed Student's *t* test. Scale bars, 100 µm. **E** Representative immunofluorescence images and quantification of BrdU-positive CMs and ECs in *ubi:loxp-eGFP-stop-loxp-npas4l* (control, n=13-14), $clo^{fv087b/+}$*; ubi:loxp-eGFP-stop-loxp-npas4l* ($clo^{fv087b/+}$, n=13-15), and $clo^{fv087b/+}$*; gata1a:Cre-ERT2; ubi:loxp-eGFP-stop-loxp-npas4l* ($clo^{fv087b/+}$; Npas4l OE, n=13-14) hearts at 7 dpa after 4-HT treatment. Arrowheads indicate proliferating CMs or ECs. Data are the mean ± SEM.; *p <0.05; ****p <0.001; one-way ANOVA with LSD test. Scale bars, 50 µm. **F** Representative immunofluorescence images of PCNA-positive CMs and ECs in control sibling+DMSO (n=15), Npas4l OE+DMSO (n=14), Npas4l OE+K02288 (n=13), and Npas4l OE+LDN193189 (n=15) hearts at 7 dpt. Arrowheads indicate proliferating CMs or ECs. Data are the mean ± SEM.; *p <0.05; **p <0.01; ****p <0.001; one-way ANOVA with LSD test. Scale bars, 50 µm.

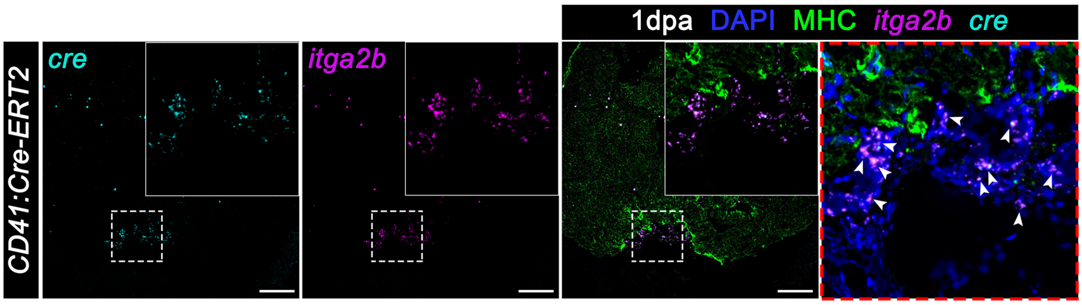


**Figure S16. CD41 promoter drives *cre* expression specifically in platelets after injury.**

Representative images of *cre-itga2b* RNAscope *in situ* hybridization co-stained MHC (labeling myocardium) on *Tg(CD41:Cre-ERT2)* heart sections at 1dpa. The insets and red-outlined panel were the high magnifications of the boxed area. Arrowheads indicate the *cre* and *itga2b* double positive cells. Scale bars, 100 µm.

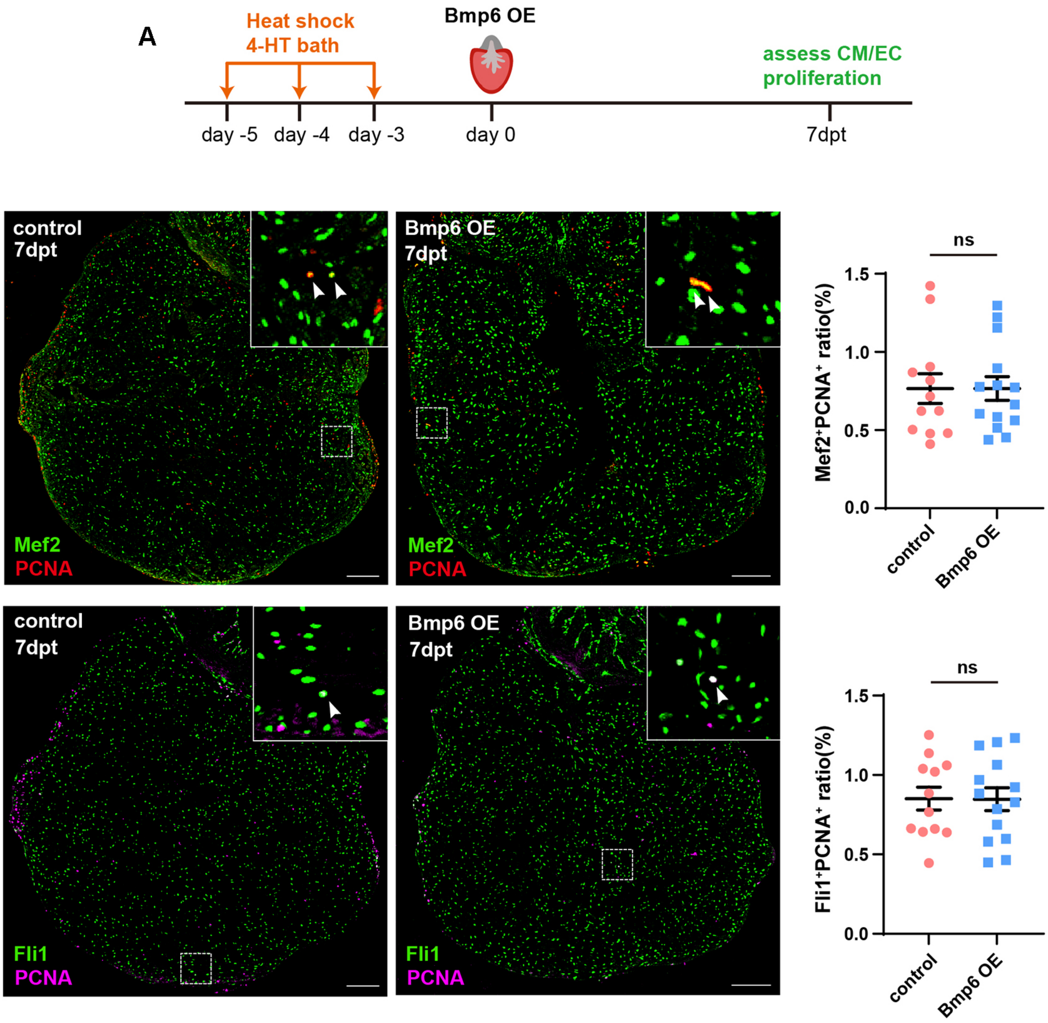


**Figure S17. Overexpression of *bmp6* has no effects on promoting uninjured CM/EC into the cell-cycle.**

**A** Schematic diagram of experimental design to explore the function of *bmp6* on promoting uninjured cardiomyocytes into the cell cycle. **B, C** Representative immunofluorescence images and quantification of PCNA-positive CMs (B) and ECs (C) in *hsp70:Cre-ERT2* (control, n=12) and *hsp70:Cre-ERT2; ubi:loxp-stop-loxp-bmp6* (Bmp6 OE, n=14) hearts at 7 days post 4-HT treatment (dpt). Insets are high magnifications. Arrowheads indicate proliferating CMs or ECs. Data are the mean ± SEM.; ns: no significance; unpaired, two-tailed Student's *t* test. Scale bars, 200 µm.

**Supplementary Tables S1-S7**

**Table S1. Marker genes for each cell type**

Gene signature and expression in each cell cluster and canonical markers used in scRNA-seq data.

**Table S2. Marker genes and GSVA for each cluster of cardiomyocytes**

Marker genes used and GSVA (Gene Set Variation Analysis) pathways enriched in cardiomyocytes subsets.

**Table S3. Marker genes and GSVA for each cluster of endothelial cells**

Marker genes used and GSVA (Gene Set Variation Analysis) pathways enriched in endothelial cells subsets.

**Table S4. Marker genes and GSVA for each cluster of platelets**

Marker genes used and GSVA (Gene Set Variation Analysis) pathways enriched in platelets subsets.

**Table S5. Npas4l CUT&TAG GO**

Gene ontology analysis of Npas4l binding genes detected by CUT&TAG. The promoter region was defined within ±10kb to TSS.

**Table S6. Npas4l binding peak motifs**

Known motif enrichment of the Npas4l binding peaks detected by CUT&TAG.

**Table S7. Bulk RNA-seq**

DEGs (Differentially Expressed Gene) of CD41$^{+}$ cells (*clo*$^{fv087b/+}$ vs. WT at 1dpa) and gata1a$^{+}$ cells (Npas4l OE vs. control at 7dpt) and corresponding pathway analysis of Npas4l-positive and -negative regulated genes from bulk RNA-seq.
